# *Candida auris* undergoes adhesin-dependent and -independent cellular aggregation

**DOI:** 10.1101/2023.04.21.537817

**Authors:** Chloe Pelletier, Sophie Shaw, Sakinah Alsayegh, Alistair J. P. Brown, Alexander Lorenz

## Abstract

*Candida auris* is a fungal pathogen of humans responsible for nosocomial infections with high mortality rates. High levels of resistance to antifungal drugs and environmental persistence mean these infections are difficult to treat and eradicate from a healthcare setting. Understanding the life cycle and the genetics of this fungus underpinning clinically relevant traits, such as antifungal resistance and virulence, is of the utmost importance to develop novel treatments and therapies. Epidemiological and genomic studies have identified five geographical clades (I-V), which display phenotypic and genomic differences. Aggregation of cells, a phenotype primarily of clade III strains, has been linked to reduced virulence in mouse and *Galleria mellonella* infection models. The aggregation phenotype has thus been associated with conferring an advantage for (skin) colonisation rather than for systemic infection. However, strains with different clade affiliations were compared to infer the effects of different morphologies on virulence. This makes it difficult to distinguish morphology-dependent causes from clade-specific or even strain-specific genetic factors. Here, we identify two different types of aggregation: one induced by antifungal treatment which is a result of a cell separation defect; and a second which is controlled by growth conditions and only occurs in strains with the ability to aggregate. The latter aggregation type depends on an ALS-family adhesin which is differentially expressed during aggregation in an aggregative *C. auris* strain. Finally, we demonstrate that macrophages cannot clear aggregates, suggesting that aggregation might after all provide a benefit during systemic infection and could facilitate long-term persistence in the host.

**Author Summary:** *Candida auris* is a single-celled fungus, a yeast, that can cause severe infections in hospital patients. This fungus is difficult to treat because it is resistant to many antifungal drugs. Therefore, to understand the processes that enhance the virulence of this yeast with a view to developing new treatments. Previous studies have found that *C. auris* can form aggregates, or clumps of cells, which may play a role in how the fungus infects people. In this study, we identified two different types of aggregation in *C. auris*, one triggered by antifungal treatment, and another controlled by growth conditions. This discovery allowed us to study aggregate formation in the same genetic background. In doing so, we found that a certain protein, an ALS-family adhesin, is involved in the aggregation process. Surprisingly, we also discovered that aggregates may promote infection by making it harder for the immune system to clear the yeast. This new understanding could help researchers develop better ways to fight *C. auris* infections.

## Introduction

The fungus *Candida auris*, first identified in 2009, has been responsible for outbreaks of infections in hospitals on six continents [1,2]. It has become a global concern due to the high levels of antifungal resistance displayed across the species, its environmental persistence, and its nosocomial transmission [3,4]. The species is divided into 5 clades which have distinct geographic origins and show different levels of intra-clade variations. The 4 main clades, clade I (South Asia), clade II (East Asia), clade III (South Africa), and clade IV (South America) are well documented, whereas only a few isolates of clade V (Iran) have been identified so far [5–7].

In fungi, morphological changes have been linked to gene expression modifications that can impact virulence and pathogenicity [8,9]. For example, the expression of adhesins and proteases are co-regulated alongside the yeast-to-hyphae transition of *Candida albicans* [10]. The yeast *C. auris* typically grows as single ellipsoidal cells, and can form filaments, but not true hyphae, under certain conditions [11,12]. Furthermore, certain *C. auris* strains can form aggregates, which are clumps of cells that cannot be dispersed by chemical or mechanical means and that are thought to be caused by a cell separation defect [13–15]. Cellular aggregation is a phenotype predominantly of clade III strains, but there are also aggregative isolates in clade I and clade II [16–18]. The biological relevance and the genetic requirements for this aggregation morphology are not fully understood yet [19], but some potential factors have recently been identified [14,17,20,21]. Indeed, this kind of cellular behaviour can also be observed in other species of the *Candida haemulonii* complex [21], to which *C. auris* belongs. *C. auris* strains displaying the aggregation phenotype seem to be less virulent, more commonly associated with skin colonisation, have greater biofilm mass, and greater environmental persistence over non-aggregative isolates [16,18,22,23]. A recent study has distinguished two types of aggregation, and has suggested that overexpression of an ALS-family adhesin, caused by copy-number variation of the corresponding locus, is involved in this [17]. So far, a major limitation of studies exploring aggregation has been that aggregative strains are often compared to non-aggregative ones from different clades.

Our study has revealed that there are two different kinds of aggregation both of which represent an inducible phenotype. One type of aggregation depends on growth conditions and can only be induced in aggregative strains (mostly clade III strains). The other depends on treatment with sub-inhibitory concentrations of echinocandins (a class of antifungal drugs) and can also be induced in non-aggregative strains. The first type of aggregation is induced by growth in rich medium and repressed in minimal medium. In contrast to media-induced aggregates which are seemingly caused by cells sticking to each other, echinocandin-induced aggregation is characterized by cells that fail to fully separate [15]. Having identified the conditions for media-induced aggregation, we exploited this to elucidate the genetic requirements for this type of aggregation. For this, we performed transcriptomic analysis to identify differentially expressed genes in aggregative and non-aggregative strains grown in rich and minimal medium. We identified a gene with homology to a *C. albicans ALS* gene that is strongly upregulated in the aggregative strain when grown in rich medium. We constructed a deletion mutant of this gene, and the resulting strain lacks the ability to aggregate in rich medium, but still aggregates in response to sub-inhibitory concentrations of echinocandins. We also show that virulence in an invertebrate infection model is affected by pre-infection culture conditions especially for strains with moderate or low virulence. If aggregation is prevented by deletion of the upregulated ALS factor in an aggregative strain, then virulence is moderately reduced. Finally, we used macrophages derived from THP-1 monocytes to demonstrate that aggregates could be difficult for the immune system to clear, and thus could potentially be responsible for persistent infections.

## Results

### Two distinct types of aggregation

The ability to aggregate (or not) has been used to classify strains of *C. auris*, and several studies have reported aggregative capacities for some clinical isolates [15,18,24]. It was thus surprising that, when grown in RPMI-1640, a defined minimal medium, we observed a lack of this phenotype regardless of clade or status as aggregative or non-aggregative strain (Fig 1A). However, we did observe that, if grown in Sabouraud dextrose broth (SabDex), an undefined rich medium, the clade III strain (UACa20) and the clade IV strain (UACa22) did form aggregates alongside single yeast cells, whereas non-aggregative clade I (UACa11) and clade II (UACa83) strains did not show any clumping of cells (Fig 1A). The inability of this clade I isolate to aggregate, as well as the aggregative ability of the clade III strain, we used in this study, agreed with previous observations [16].

**Fig 1.**
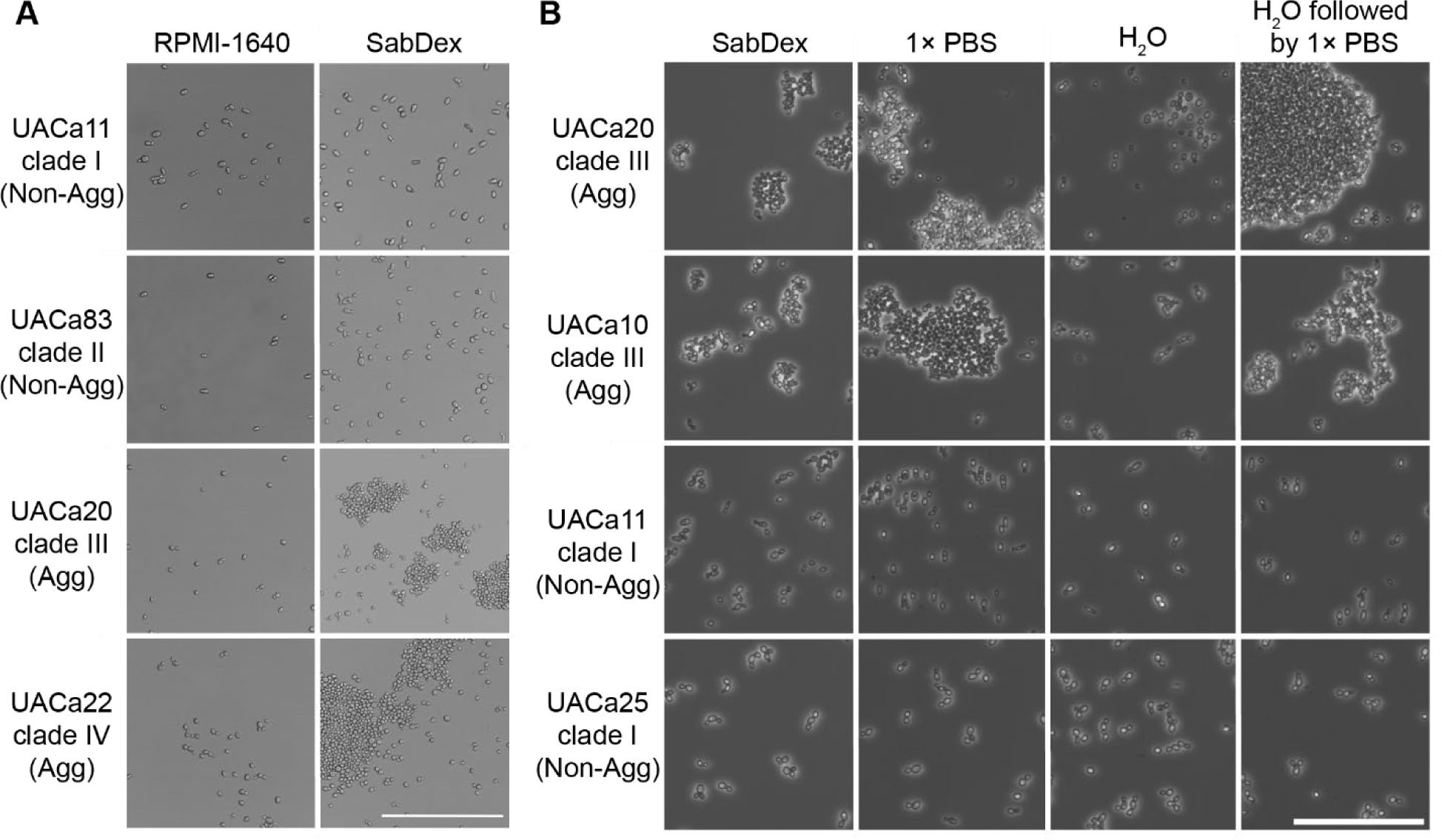
Aggregation is an inducible phenotype. (**A**) Changes in growth media induce aggregation only in aggregative strains. Light microscopic (brightfield) images of cells of the indicated strains (left) in the indicated medium (top). Cells grown in RPMI-1640 displayed no aggregation regardless of ability to aggregate. Only aggregative (Agg) strains grew as aggregates when grown in SabDex while non-aggregative (Non-Agg) strains remained as single cells. (**B**) Aggregation is lost when media is replaced with water but is retained when replaced with 1× PBS. Light microscopy (phase contrast) of cells grown overnight in SabDex followed by either replacement of media with ddH2O, 1× PBS, or ddH2O followed by 1× PBS. Non-aggregative (Non-Agg) strains were unaffected by the suspension liquid while aggregative (Agg) strains retained aggregation when suspended in 1× PBS, but aggregates were noticeably reduced after washing with ddH2O. Scale bars represent 50 µm.

We also found that, after growth in SabDex, aggregation was affected by the chemical composition of the liquid used to resuspend the cells. Aggregating cultures retained aggregation when resuspended in 1× PBS (phosphate-buffered saline), but aggregation was greatly reduced when cells were resuspended in ddH2O (Fig 1B). Interestingly, cells that were dis-aggregated in ddH2O, immediately re-aggregated when they were then resuspended in 1× PBS (Fig 1B). Non-aggregative strains remained as single cells regardless of suspension liquid (Fig 1B). This strongly indicates that a cell wall component that remains present regardless of the change in suspension liquid is associated with aggregation.

It has been reported that *C. auris* strains aggregate in response to sub-inhibitory concentrations of two classes of antifungals, azoles, and echinocandins [15,16]. We focused on two echinocandins, caspofungin (CSP) and micafungin (MFG) as these drugs are used preferentially in the clinical setting to treat *C. auris* infections (<u>CDC Guidelines</u>, <u>PHE Guidlines</u>). E-test strips were used to determine minimum inhibitory concentrations (MICs) for the strains tested (Table S1). Growth was not completely abolished at any concentration of CSP but was clearly diminished above a certain concentration which we recorded as the MIC90 value (Table S1, Fig S1). Growth in the presence of MFG was abolished above 0.094 mg/L for clade I strains UACa11 and UACa25 (Table S1, Fig S2). However, for the clade III strains (UACa10 and UACa20) MIC90s of 0.094 mg/L and 0.064 mg/L, respectively, were determined as residual background growth was observed above these concentrations (Table S1, Fig S2). Therefore, 32 mg/L CSP and 0.075 mg/L MFG were chosen as subinhibitory concentrations.

Echinocandin-induced aggregation experiments were performed in RPMI-1640 to avoid clade III strains forming media-induced aggregates. Indeed, both the aggregative clade III strain (UACa20) and the non-aggregative clade I strain (UACa11) formed aggregates when grown in RPMI-1640 containing either 32mg/L CSP or 0.075 mg/L MFG (Fig 2A-B), but not in the presence of the DMSO vehicle (Fig S3). These aggregates did not disperse when cells were resuspended in ddH2O, indicating that antifungal-induced aggregation differs from media-induced aggregation (Figs 1B, 2A-B). This difference was also noticeable when looking at the liquid cultures. After 5 minutes without agitation there was noticeable sedimentation of the media-induced aggregates in the aggregative UACa20 (clade III) strain not seen in any of the other cultures (Fig 2C). However, both the non-aggregative UACa11 (clade I) isolate and the aggregative UACa20 (clade III) strain largely remained in suspension when echinocandin-dependent aggregates were formed (Fig 2C). Neither media-nor echinocandin-induced aggregation was reflected in an altered colony morphology on solid medium (Fig 2D).

**Fig 2.**
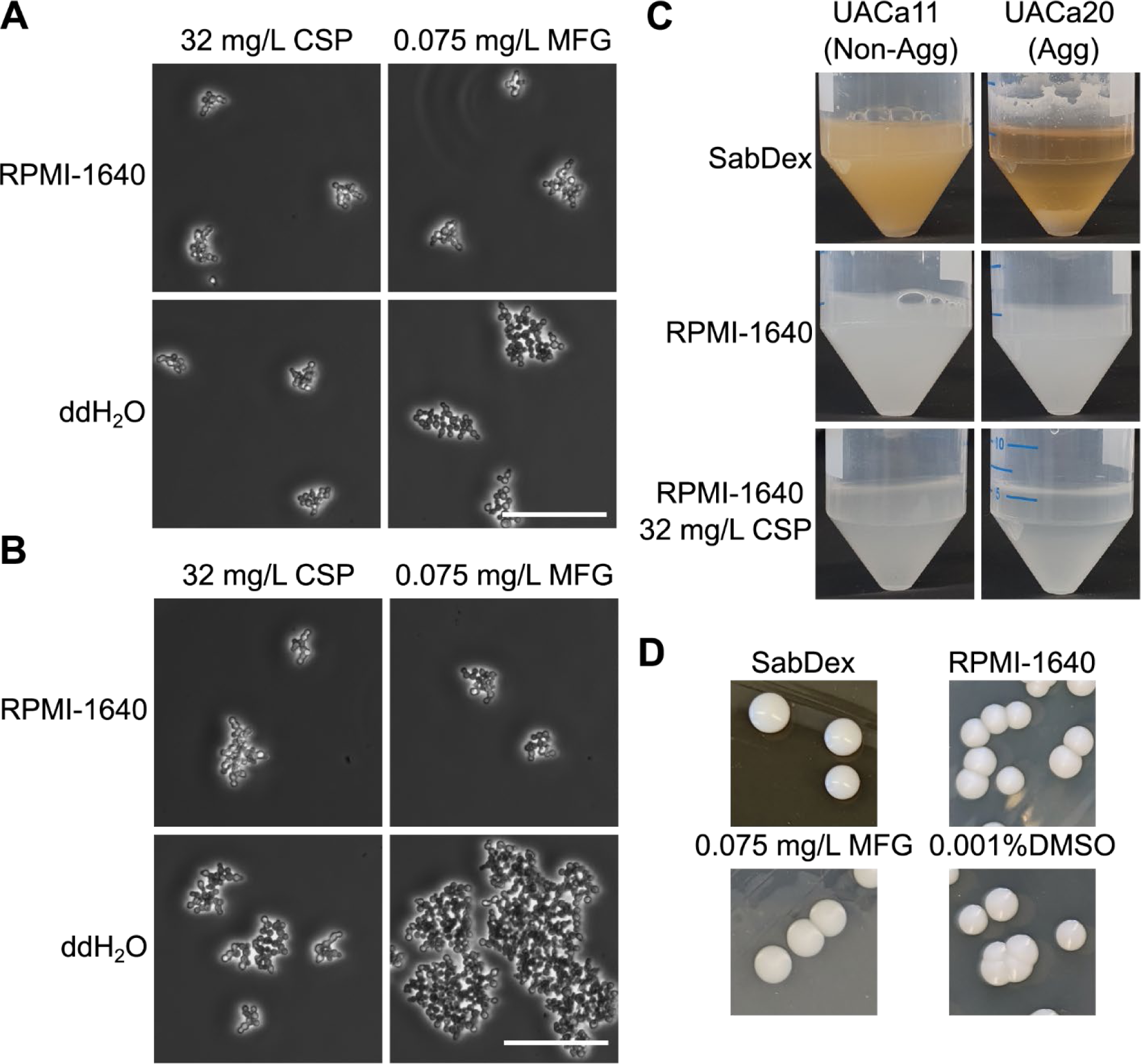
Sub-inhibitory concentrations of echinocandins induce a different type of aggregation (clustering). (**A**) UACa20 (clade III) and (**B**) UACa11 (clade I) grown overnight in RPMI-1640 containing either 32 mg/L CSP or 0.075 mg/L MFG, images were taken with a light microscope (phase contrast) of cells in media and when resuspended in ddH2O, showing drug-induced clusters remain intact after washing with ddH2O. Scale bars in (A) and (B) represent 50 µm. (**C**) Differences in sedimentation were not observed in the RPMI-1640 cultures or the drug-induced clusters, while media-induced aggregates did fall out of suspension. (**D**) Colony morphology was not conspicuously impacted by aggregation.

The capacity of media-induced aggregates to dissociate and reform depending on the suspension liquid, suggests the involvement of a component of (or associated with) the cell wall, whereas the stability of antifungal-induced aggregates is consistent with a cell separation defect [15]. To explore these phenotypes in more detail, we used calcofluor white (CFW) to stain total cell wall chitin on paraformaldehyde-fixed cells. Media-induced aggregates contained a mix of larger rounded cells marked with multiple bud scars alongside smaller ellipsoidal cells (Fig 3A). It was also noted that the media-induced aggregates started to fall apart when proteinase K treatment was used during preparation, causing aggregates to spread out across glass slides during preparation for microscopy. No pressure was applied during preparation as a hard-set mounting medium was used. In contrast, echinocandin-induced aggregates appeared to grow from a small cluster of cells with daughter cells emanating from a central point (Fig 3B). These tight clusters of cells retained their shape during microscopy even after treatment with proteinase K. The daughter cells appeared to remain attached to their mothers indicating a cytokinesis defect similar to rapamycin-treated cells [25]. A lack of obvious bud scars on cells at the edge of clusters corroborates this interpretation (Fig 3B) [25].

**Fig 3.**
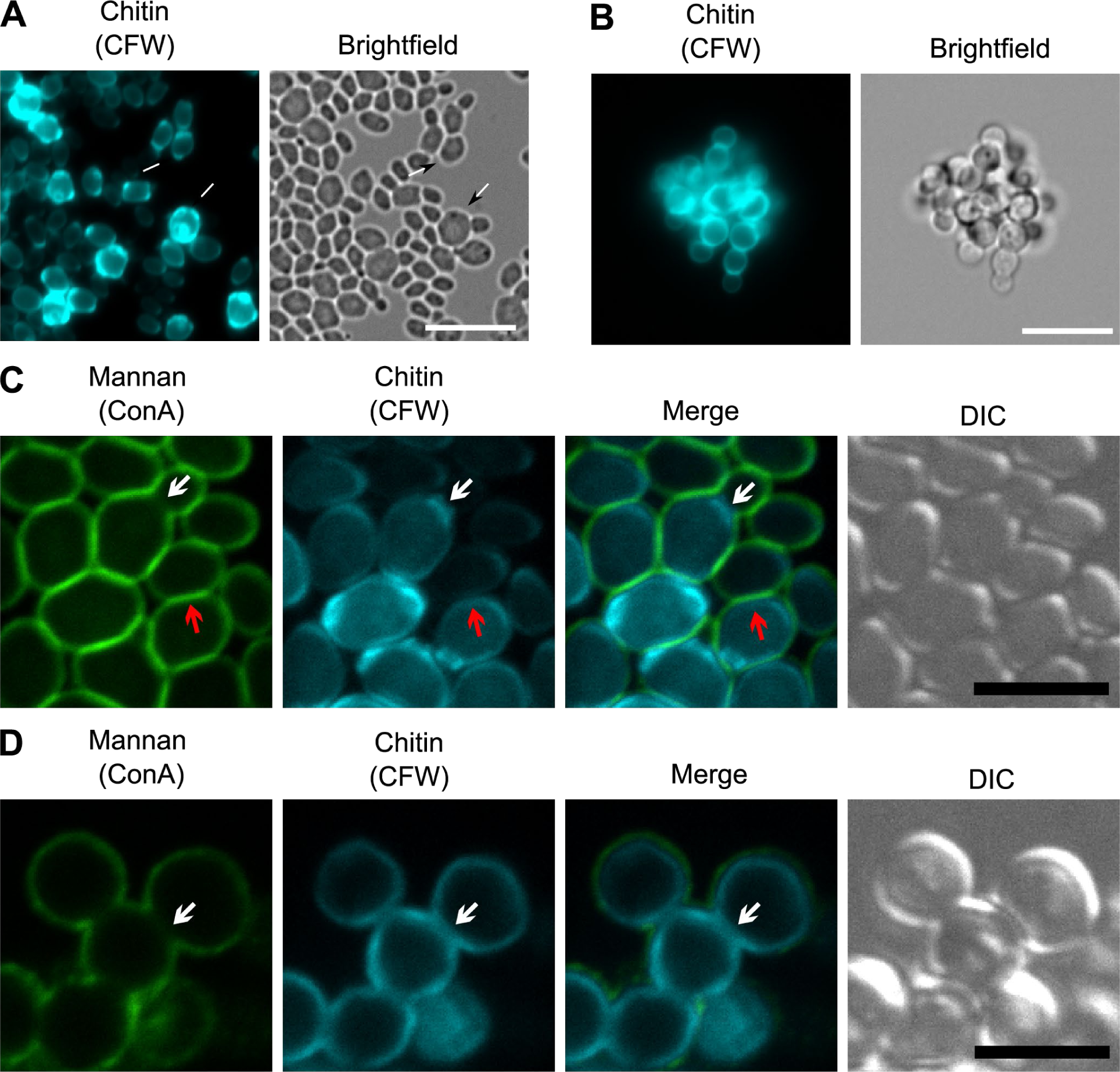
Differences in chitin bud scars and the coverage of outer cell wall mannan indicate differences between aggregation types. (**A, B**) UACa20 (clade III) cells grown overnight in SabDex (A) or RPMI-1640 containing 32 mg/L CSP (B), cell wall chitin is visualized with 10 µg/mL CFW. White arrows point to bud scars without daughter cells grown in SabDex indicating full cell separation after division, this is not seen on antifungal-induced aggregation. Scale bars in (A) and (B) represents 10 µm. (**C, D**) Cell wall mannan and chitin stained with ConA and CFW, respectively, visualized by confocal microscopy to generate image sections of entire aggregates. (**C**) Cells grown in SabDex are completely outlined by mannan and chitin staining (red arrows), even where cells are closely juxtaposed, only actively dividing cells with small daughter buds appear to have a break in the mannan outer cell wall layer (white arrows). (**D**) When grown in RPMI-1640 containing 0.075 mg/L MFG cells are completely bounded by chitin, but display a lack of mannan staining at cell-cell junctions (white arrows) indicating a cell separation defect. In (C) and (D) the scale bars represent 5 µm.

To better understand whether antifungal-induced aggregation was due to a cell separation defect, confocal microscopy was used to visualize aggregates in three dimensions. UACa20 (clade III) cells grown in SabDex were double-stained with CFW (chitin) and Concanavalin A (ConA) (cell wall mannan). Media-induced aggregates displayed an intact chitin cell wall that was completely covered by mannan (Fig 3C, red arrows). Only small budding cells lacked the mannan outer cell wall layer between mother and daughter cells (Fig 3C, white arrows). In contrast, cells in antifungal-induced aggregates had a chitin layer completely surrounding them, but there was a lack of a mannan layer where cells were closely juxtaposed (Fig 3D, white arrows), similar to small budding cells in media-induced aggregates. Again, this is consistent with the idea that antifungal-induced aggregation is the result of a cell separation defect, as previously suggested [13–15,17].

During the preparation of cells for microscopy we noted that there was a lack of cohesion of media-induced aggregates as well as loss of aggregation after treatment with proteinase K for 1 hour at 50 °C. This was never observed for echinocandin-induced aggregates. To quantify this observation, aggregates were counted using a haemocytometer, and 1 × 10^6^ aggregates were heated at 50 °C for 1 hour with or without 12.5 µg of proteinase K. The log2-fold change of aggregates remaining after treatment was determined relative to number of aggregates before treatment. As predicted, the number of media-induced aggregates showed a significant decrease when proteinase K was present. For UACa20 (clade III), there was a significant ∼15-fold decrease (*p* = 0.03, independent samples t-test), and UACa10 (clade III) showed a significant ∼28-fold decrease (*p* < 0.001, independent samples t-test) in aggregates after treatment with proteinase K (Fig 4A). These results are consistent with observations by others [17,21], and suggest that the cell wall component involved in media-induced aggregation is not perturbed by temperatures up to 50 °C. Neither CSP- nor MFG-induced aggregates were significantly reduced by proteinase K treatment (Fig 4B-C).

**Fig 4.**
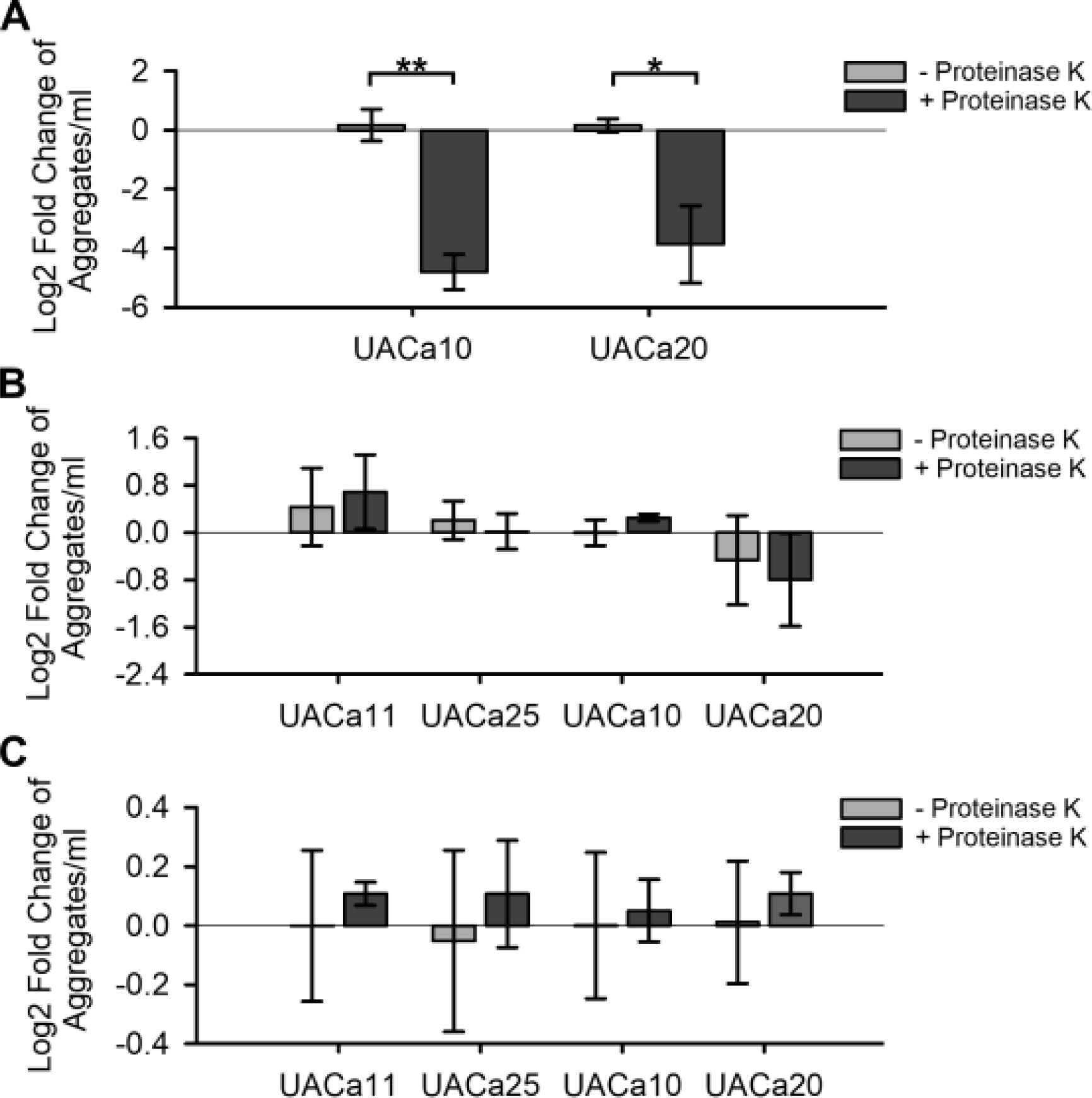
Proteinase K treatment significantly reduces media-induced aggregation but not antifungal-induced clustering. Log2-fold change in aggregate/cluster numbers grown in SabDex (**A**), 32 mg/L CSP (**B**), or 0.075 mg/L MFG (**C**) after incubation at 50 °C for 1 hour with or without 12.5 µg of proteinase K. \**p* < 0.05, \*\**p* < 0.001, as determined by independent samples t-test.

Based on these observations, to semantically distinguish these phenotypes, we propose that media-induced aggregation is henceforth referred to as “aggregation”, whereas aggregation caused by a cell separation defect is called “clustering”. We focused on aggregation induced by growth conditions to understand why this phenotype only occurs in some isolates of *C. auris* and what the biological relevance of this phenotype might be.

### Media-induced aggregation is not correlated with cell wall ultrastructure

Dispersion of aggregates by proteinase K suggested that a proteinaceous cell wall component is involved in media-induced aggregation. To rule out other cell wall ultrastructure changes and a potentially obscure cell separation defect, transmission electron microscopy (TEM) was performed. Cells were grown in either RPMI-1640 or SabDex before high-pressure freezing fixation without washing. No noticeable differences in the cell wall ultra-structure were observed between conditions for all strains regardless of the growth conditions (Fig 5A).

**Fig 5.**
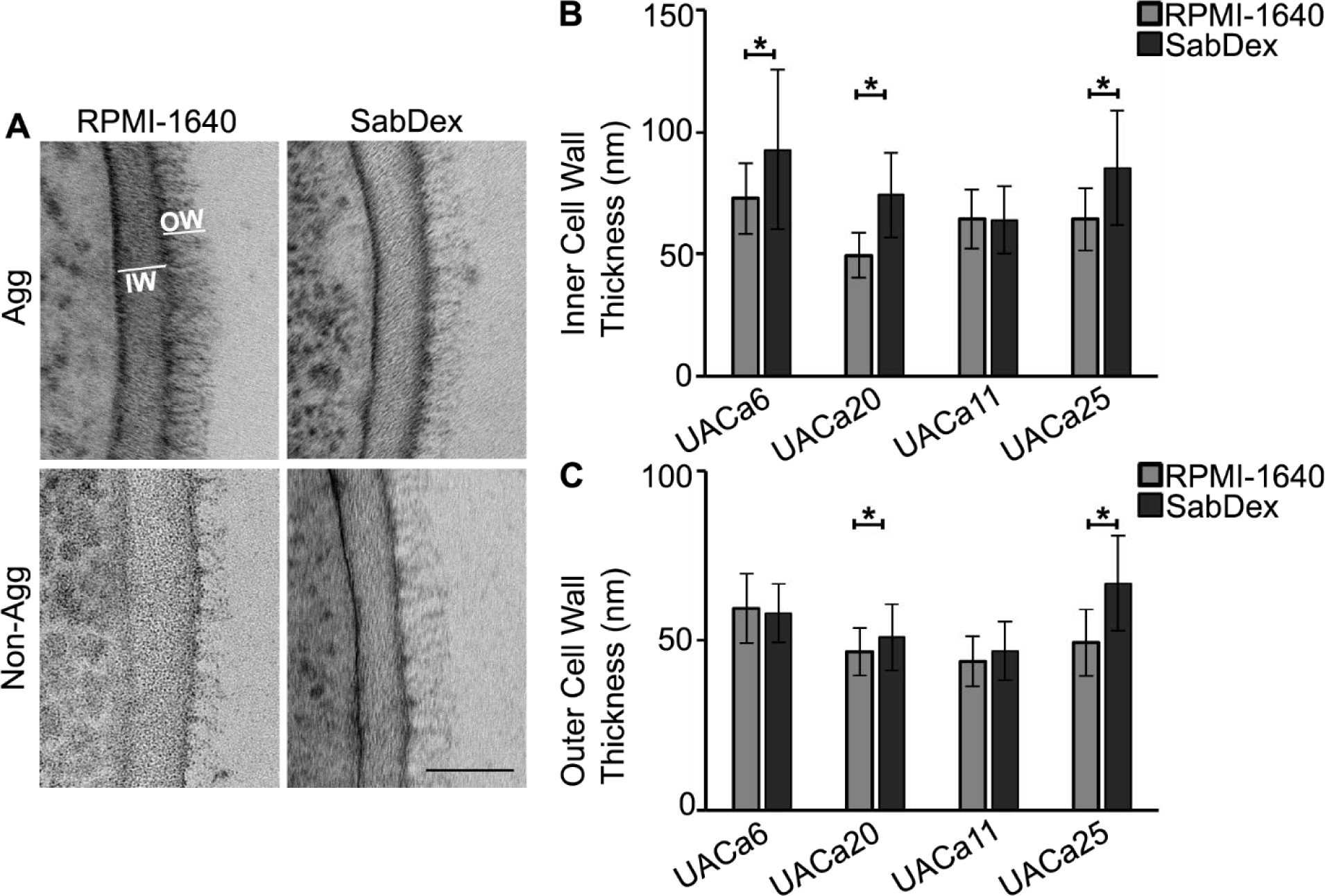
Media-induced aggregation does not cause changes to the cell wall ultrastructure. Cells grown in indicated media were fixed by high pressure freezing for transmission electron microscopy TEM. (**A**) Representative images from TEM of cell wall of an aggregative (Agg) and a non-aggregative (Non-Agg) strain, outer cell wall (OW) and inner cell wall (IW) are indicated. Scale bar represents 50 nm. (**B**) Inner cell wall measurements show strain-specific changes in response to culture conditions. (**C**) Length of mannan fibrils (outer cell wall) shows moderate changes in response to culture conditions in two isolates, UACa20 (clade III) (Agg) and UACa25 (clade I) (Non-Agg). \**p* < 0.05 as determined by independent samples t-test.

The diameters of the inner and outer cell walls for 30 cells from each condition were measured. The diameter of the inner cell wall of non-aggregative strain UACa25 (clade I) showed a significant increase of 21 nm (*p* < 0.001, independent samples t-test) when grown in SabDex compared to RPMI-1640, but there was no significant difference for UACa11 (clade I), a second non-aggregative strain. Aggregative clade III strains UACa6 and UACa20 grown in SabDex had significantly thicker inner cell walls by 20 nm (*p* < 0.001, independent samples t-test) and 25 nm (*p* < 0.001 independent samples t-test), respectively, than when grown in RPMI-1640 (Fig 5A). The mannan fibrils (outer cell wall) of UACa25 (clade I) were significantly longer by 18 nm (*p* < 0.001, independent samples t-test) when grown in SabDex compared to RPMI-1640 (Fig 5B). While UACa20 (clade III) grown in SabDex had slightly longer mannan fibrils by 4 nm (*p* = 0.029, independent samples t-test) compared to when grown RPMI-1640. There were no significant differences of outer cell wall diameters for UACa6 (clade III) and UACa11 (clade I). Although there were differences in inner and outer cell wall thickness between the two growth conditions, there was no clear trend which separated aggregative from non-aggregative isolates.

### Transcriptomic analysis identifies clade-specific and aggregation-specific differentially expressed genes

We had shown that the aggregation phenotype is inducible in aggregative isolates, which mostly belong to clade III (Fig 1A). Therefore, we opted to exploit this by using RNA-seq to characterize genetic requirements and identify drivers of aggregation in UACa20 (clade III) compared to the non-aggregative clade I strain UACa11. We compared differentially expressed genes (DEGs) between growth in RPMI-1640 and SabDex for both strains, and then compared these DEGs between UACa20 (clade III) and UACa11 (clade I) to identify changes in transcription that may be important for aggregation. Differential expression analysis was carried out by pairwise comparison between each sample group each consisting of three independent samples, which allowed identification of DEGs between growth in SabDex compared to RPMI-1640 with genes being identified from the *C. auris* B11221 genome (UACa20 in this study). For UACa11 (clade I), from a total of 5,419 candidate open reading frames 3,191 significant DEGs at FDR < 0.05 were identified (Fig 6A). For UACa20 (clade III), from a total of 5,431 candidates 3,393 significant DEGs at FDR < 0.05 were determined (Fig 6B). There were 997 common DEGs between the two strains. These genes may be involved in adapting to growth under the two different media conditions (Fig 6C), and hence were excluded from further analysis. Genes with a log2FC difference greater than 2 between UACa20 (clade III) and UACa11 (clade I) but with increased expression in SabDex in both strains were identified (Table S2). Furthermore, we also extracted genes that displayed expression in opposite directions between UACa11 (clade I) and UACa20 (clade III) when grown in SabDex (Table S3). Ultimately, we decided to focus on genes that were uniquely upregulated in the aggregative UACa20 strain compared to the non-aggregative UACa11 strain during growth in SabDex, as we hypothesized that this would lead us to a positive effector of aggregation. UACa11 (clade I) had 505 unique DEGs during growth in SabDex, while UACa20 (clade III) had 658 unique DEGs (Fig 6C).

**Fig 6.**
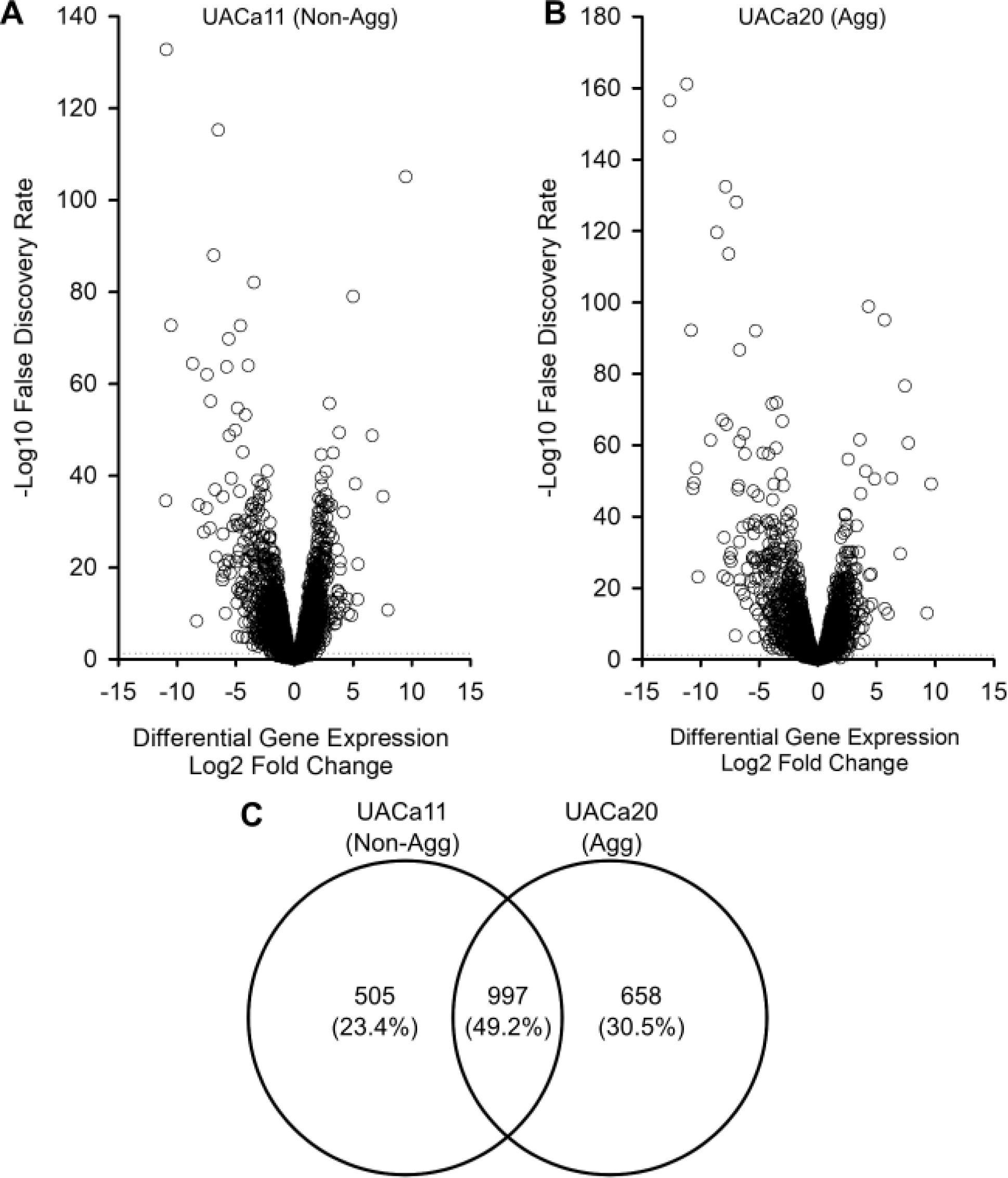
Differential expression of genes (DEGs) in an aggregative and a non-aggregative strain grown in SabDex or RPMI-1640. Aggregative (Agg) clade III strain UACa20 and non-aggregative (Non-Agg) clade I strain UACa11 were grown in the specified media for 16 hours and RNA was extracted for RNA-seq, this was repeated three times on independent colonies at different times. (**A, B**) Volcano plots of the total DEGs for UACa11 (A) and UACa20 (B) are shown with the dotted line indicating an FDR = 0.05. (**C**) Venn diagram of statistically significant (FDR < 0.05) DEGs, numbers indicate how many DEGs are unique to UACa11 and UACa20 and what portion of DEGs are shared between the strains.

The top 10 unique genes upregulated during SabDex growth for UACa11 (clade I) and UACa20 (clade III) are shown in Tables S4 and S5, respectively. To understand their possible gene functions the *C. albicans* homologues were identified. First, the B11221 nucleotide sequence of each of those genes (Tables S4-S5) was BLAST-searched against the B8441 genome and the corresponding gene/systematic ID names in B8441 were obtained. The B8441 gene identifiers were then used in the batch-download feature of the *Candida* genome database (http://www.candidagenome.org) to identify orthologues and best hits in the *C. albicans* SC5314 genome. The best hit in UACa11 (clade I) (Table S4) and the second-best hit in UACa20 (clade III) (Table S4) have been identified as homologous to *C. albicans ALS4*. However, these represent two different *C. auris* genes and they show differential regulation in regard to growth media, which was unexpected. ALS (Agglutinin-Like Sequence)-containing proteins form a family of nine adhesins in *C. albicans* with various roles including adherence to plastic and host tissues [26]. *C. auris* genomes seem to contain three separate loci that harbour an Agglutinin-Like Sequence (ALS) [27,28]. Representatives from clade II miss one of the *ALS*-family genes, and in some strains a particular *ALS* gene might be amplified (copy-number variation) [17,28]. Intriguingly, one of these factors (B9J08_004112, XP_028889036) has mostly been studied, and varyingly touted as the ortholog to *C. albicans* Als3 [29], Als4 [17,28], or Als5 [22]. This is confusing, and as ALS factors in *C. albicans* have distinct roles and different expression patterns, it is imprudent to assign orthology in *C. auris* when function and gene expression cues have not been established in this species. Indeed, phylogenetic analysis indicates that ALS-family members cluster by species and not by a particular type of ALS from different species [28], this also seems to be the case for Hyr/Iff-like adhesins [30]. To prevent further confusion, we follow the solution proposed by O’Meara and colleagues [31] to use the unique number in the systematic ID as gene number. And therefore, B9J08_004112/XP_028889036 should be called Agglutinin-<u>L</u>ike <u>S</u>equence 4112 (*ALS4112*/Als4112), B9J08_002582 *ALS2582*, and B9J08_004498*ALS4498*. This nomenclature should avoid potential confusion with ALS-type proteins from other species where exact relationships are hard to establish.

### Als4112 is necessary for aggregation

The *C. albicans* ALS-family proteins are found on the cell surface, play prominent roles in adhesion to host tissues and other surfaces, and are important for virulence [26,32,33]. As media-induced aggregation is likely caused by a cell surface protein and *ALS4112* displays differential expression during SabDex and aggregative growth in UACa20 (clade III), this gene was taken forward as a candidate for potentially playing a role in aggregation. We attempted to generate a clean *als4112* deletion in the aggregative strain UACa20 (clade III). Unfortunately, due to the difficulties in the genetic manipulation of *C. auris* [12,19,34], the one mutant strain we obtained harboured an approximate 21.3 kb deletion encompassing the *ALS4112* locus and genes CJI97_004175, CJI97_004176, CJI97_004177, CJI97_004178, CJI97_004179, CJI97_004180, as determined by inverse PCR (Fig S4). We proceeded with this mutant as the additional genes deleted were not present as DEGs in our RNA-seq dataset and are, therefore, not thought to play a role in aggregation; the *als4112* mutant strain also did not display an obvious growth defect. This *als4112* mutant failed to aggregate after 24 hours of growth in SabDex but did still cluster upon exposure to 0.075 mg/L MFG (Fig 7A). This reinforces the idea that there are two distinct types of aggregation, because the same type of cell separation defect is observed in echinocandin-induced clustering in the wild-type clade III UACa20 strain, as well as in the *als4112* mutant made in the same background (Figs 3D and 7B). Thus, *ALS4112* appears to be a key genetic requirement for media-induced aggregation but not for echinocandin-mediated clustering.

**Fig 7.**
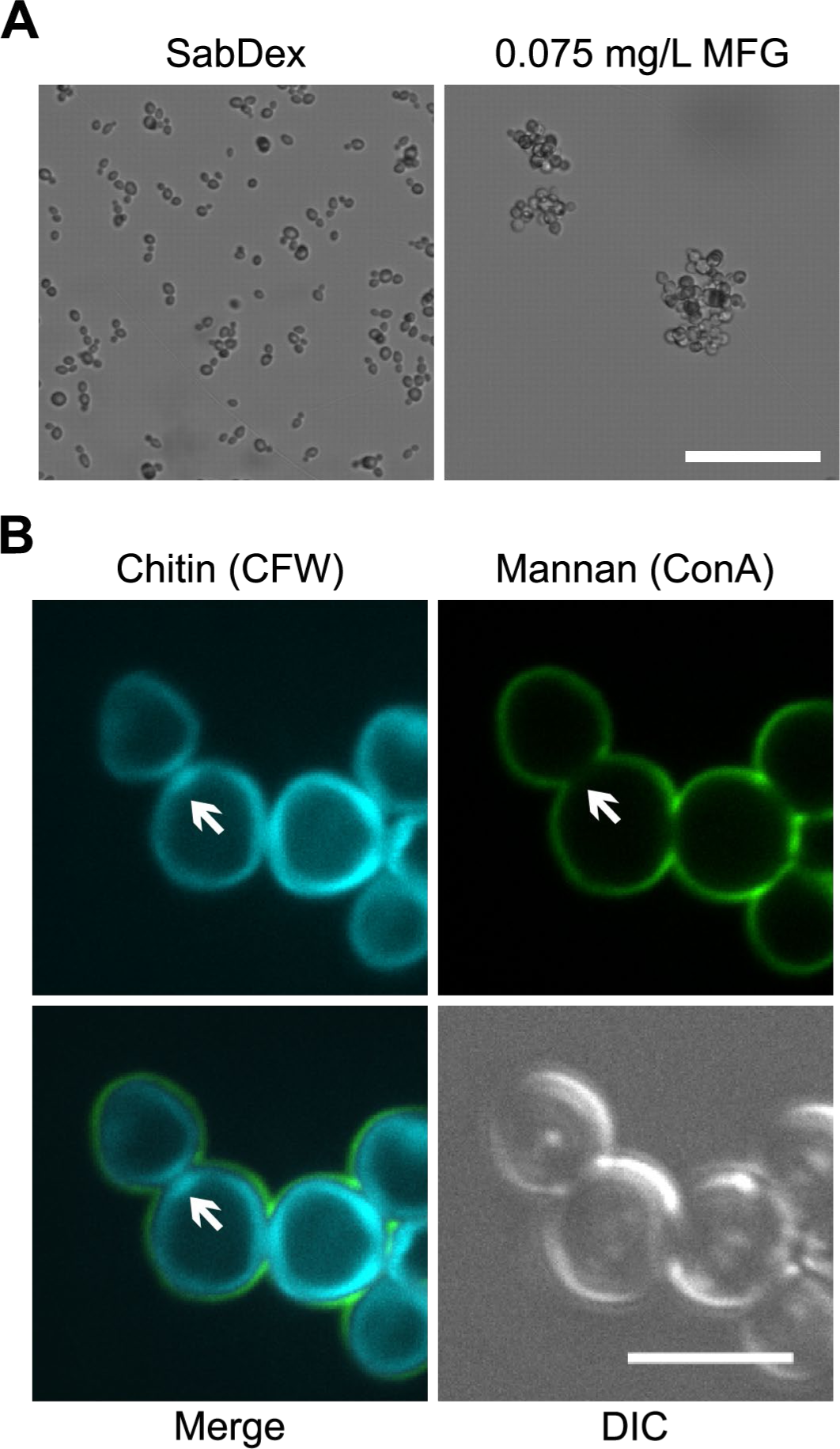
*ALS4112* seems to be required for media-induced aggregation, but not for antifungal-induced clustering. (**A**) Light microscopy images (contrast) of the *als4112*Δ mutant grown SabDex or RPMI-1640 containing 0.075 mg/L MFG for 24 hours. Cells grown in SabDex failed to aggregate, whereas cells grown in the presence of MFG did form clusters. Scale bar represent 10 µm. (**B**) The *als4112*Δ mutant was grown for 24 hours in RPMI-1640 containing 0.075 mg/L MFG and was imaged by confocal microscopy. Total cell wall chitin was stained with CFW and mannans stained with ConA. White arrows highlight features consistent with antifungal-induced clustering, where cells are completely surrounded by chitin staining, but display a lack of mannan staining at cell-cell junctions. Scale bar represents 5 µm.

### Virulence depends on growth conditions of cells before inoculation

There are mixed reports regarding the impact of aggregation on virulence in *Galleria mellonella* and mouse infection models, with some studies showing reduced virulence of aggregative strains in contrast to others observing no such a correlation [13,18,35]. Indeed, an infection study conducted in neutropenic mice found that there was no obvious association of a given strain’s virulence with either its aggregative capacity or it being isolated from the blood stream [18]. Given that in other *Candida* species, such as *C. albicans*, virulence can be influenced by cellular morphology [10], we decided to investigate whether virulence was impacted by the aggregation morphology or was isolate-dependent using an *in vivo G. mellonella* infection assay. Fungal cells were grown in either RPMI-1640 as single cells or pre-grown in SabDex inducing the aggregation phenotype in aggregative isolates UACa20 (clade III), UACa6 (clade III), and UACa23 (clade IV) while non-aggregative isolates UACa11 (clade I), UACa25 (clade I), and UACa83 (clade II) remained as single cells, before being injected into the moth larvae.

All isolates, with the exception of UACa25 (clade I), showed a significant (*p* < 0.05) increase in killing regardless of their ability to aggregate when grown in SabDex, rather than RPMI-1640, before inoculation (Fig 8A-F). This suggests that virulence does not depend directly on the ability to aggregate, but is more likely driven by more complex strain-specific traits influenced by pre-inoculation culture conditions. Being grown in rich medium (SabDex) before inoculation in particular boosts pathogenicity of strains with low to moderate virulence (Fig 8C-F), whereas the two clade I strains receive comparatively little benefit (Fig 8A-B). Indeed, also in *C. albicans* growth conditions influence virulence in invertebrate and vertebrate models of infection [36,37].

**Fig 8.**
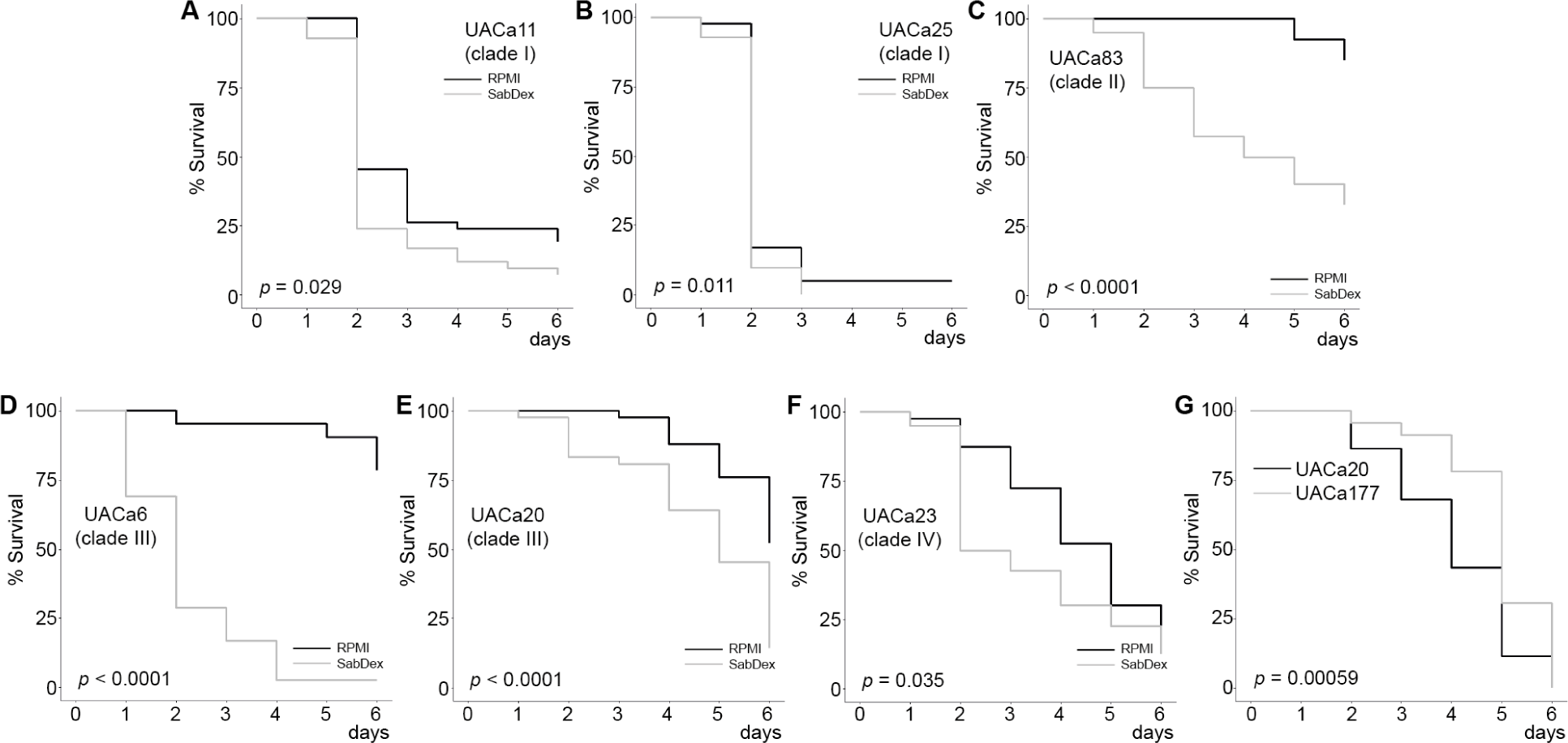
Virulence in the *G. mellonella* infection model is dependent on pre-growth media and Als4112. (**A-C**) Non-aggregative and (**D-F**) aggregative *C. auris* strains were grown in RPMI- 1640 or SabDex before inoculation into *G. mellonella* larvae. All strains except UACa25 (clade I) in (B) showed significant differences in virulence dependent on the media used to grow the fungal cells before inoculation. (**G**) Wild-type (UACa20) and *als4112*Δ (UACa177) *C. auris* cells were grown in SabDex before inoculation into *G. mellonella* larvae (A log-rank test was used to calculate *p*-values. 30 larvae in 3 independent experiments were used to assess virulence.

As the *als4112* deletion strain in the UACa20 background allowed us to directly assess whether aggregation impacts virulence. We also compared the wild-type UACa20 isolate with UACa177 in the *Galleria* infection model after growing these strains in SabDex, where UACa20 aggregates and UACa177 does not. The aggregating wild-type UACa20 isolate was moderately, but significantly, more virulent than the non-aggregating *als4112*Δ mutant strain UACa177 (Figs 8G). This suggests that ALS-dependent aggregation has a positive effect on virulence rather than a negative one as previously thought [13].

### Aggregates are not cleared by macrophages

Aggregates have been reported in tissue from sacrificed animals [18]. Given the bulky nature of aggregates, we wanted to understand how they would interact with macrophages. THP-1 monocytes were differentiated into macrophages with 200 nM phorbol 12-myristate 13-acetate (PMA) and co-cultured at a multiplicity of infection (MOI) of 1:3 with UACa20 (clade III) cells, which were either grown in SabDex, RPMI-1640, RPMI-1640 containing 0.001% DMSO, or RPMI-1640 containing 0.075 mg/L MFG and 0.001% DMSO. Cells were washed and added to CO2-independent medium for co-culture. Aggregates were counted as one unit for MOI; because cell uptake by macrophages was not quantified, we do not consider this an issue. For cells grown in RPMI-1640 or RPMI-1640 containing 0.001% DMSO, there was no noticeable defect in uptake of single yeast cells (Videos V1 & V2). When presented with media-induced aggregates (Fig 9A, Video V3) or MFG induced clusters (Fig 9B, Video V4) macrophages struggled to engulf the mass of fungal cells with fungal growth occurring at the site not occupied by the macrophage. Media-induced aggregation is not complete, as it produces a heterogenous population of single cells and large aggregates. The single yeast cells under these co-culture conditions are readily taken up by macrophages (Video V5). This indicates that there is no issue with sensing the *C. auris* cells growing under aggregating conditions but that the ability of macrophages to clear the fungal aggregates is dampened. Aggregation and clustering could potentially be causes for long-term persistence of fungal cells in the host and thereby contribute to increased virulence.

**Fig 9.**
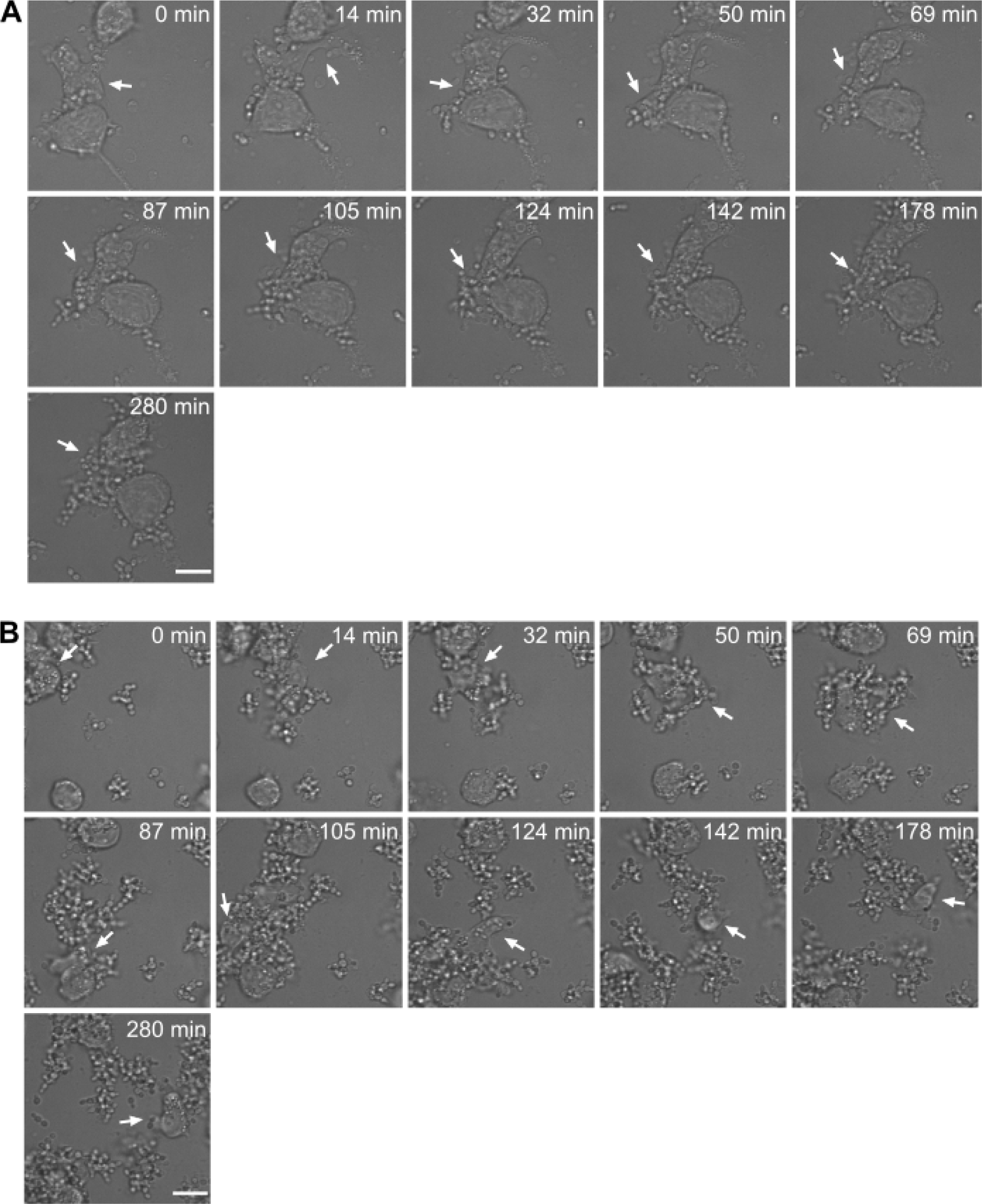
Macrophages have difficulty in clearing large aggregates and clusters. (**A**) Aggregates of strain UACa20 (clade III) grown in SabDex were co-incubated with THP-1 derived macrophages, and one macrophage was followed for the duration of the experiment (white arrow), taken from Video V3. (**B**) Clusters of strain UACa20 grown in RPMI-1640 containing 0.075 mg/L MFG, one macrophage was followed for the duration of the experiment (white arrow), taken from Video V4. Scale bars represent 15 µm.

## Discussion

Here, we show that aggregative *C. auris* strains, usually belonging to clade III, only aggregate when grown in a rich medium, such as SabDex or YPD (Fig 1A). This aggregation phenotype is repressed in a minimal medium, such as RPMI-1640, in which aggregative strains grow as single-celled yeasts, like non-aggregative strains (Fig 1A). A second type of aggregation, which we suggest calling “clustering”, was seen when cells were exposed to sub-inhibitory concentrations of echinocandins. This clustering can be induced in aggregative and non-aggregative strains (Fig 2). Such changes in morphological phenotype in *C. auris,* which are triggered by environmental cues, are likely biologically relevant. Similar morphological transitions are observed in other *Candida* species where the switches are often associated with cell stress, such as increased temperature or lack of nutrients [38,39]. SabDex is a rich undefined medium, so the switch of aggregative strains to grow as aggregates suggests that either rich medium is causing some cellular stress or contains specific chemical triggers for such a switch. Indeed, our work highlights the importance of selecting appropriate growth conditions for experimentation with *C. auris,* especially when the aggregation phenotype is to be dissected. Hence, publicly available data from previous work should be reanalysed cautiously [15], as effects of media-induced aggregation and antifungal-induced clustering could potentially confound each other.

It was originally assumed that aggregates in *C. auris* formed due to a lack of cell separation [13]. Indeed, some genetic requirements for this type of cell clustering have been identified by elegant forward genetic screening [14,40]. We only saw this particular phenotype when we exposed *C. auris* strains to echinocandins, and this did not happen when *C. auris* clade III strains were grown in rich media (Fig 1A & 3). However, Bing *et al*. (2023) characterized a clinical *C. auris* clade I isolate that forms clusters via a cell separation defect, which seems to be inherent to this particular strain [17]. It will be interesting to assess whether clustering in this strain can be modulated by growth condition or is exacerbated by treatment with echinocandins. In the same study [17], a *C. auris* clade III strain was shown to undergo proteinase-sensitive aggregation which is associated with overexpression of an ALS-type adhesin (B9J08_004112). For reasons outlined above, we refer to B9J08_004112 as *ALS4112*. Overexpression of Als4112 (Bing *et al*. call it Als4 [17]) in some clinical *C.auris* clade III isolates is apparently caused by copy-number variation of that gene apparently linked to adherence and biofilm forming capability [17]. Copy-number variation and induced gene expression levels also neatly explain our observation that a change in media provokes aggregation. In such a scenario, an aggregative strain expresses Als4112 at low levels when grown in minimal medium (RPMI-1640) but has a high-enough copy number of *ALS4112* [17] to produce sufficient protein molecules to enable aggregation once expression is induced. Here, we find that the non-aggregative UACa11 clade I strain does not upregulate expression of *ALS4112* when grown in SabDex but a different *ALS* gene, *ALS2582* (B9J08_002582). Als2582 at 861 amino acids is considerably smaller than Als4112 (1804 amino acids). Our data suggest that Als2582 does not drive aggregation, as UACa11 (clade I) does not form aggregates. Whether this particular adhesin is involved in biofilm formation, adherence to host cells, or adherence to other surfaces (medical equipment) remains to be tested. Indeed, a recently identified *C. auris*-specific adhesin, Scf1, contributes to virulence by promoting adhesion to both biological and abiotic surfaces [31].

We show that antifungal treatment causes a cell separation defect which leads to the formation of cell clusters, and that this phenotype is independent of the ability to aggregate in response to the growth media, which has been assumed to be the typical aggregation phenotype [13,17]. Although this is an important distinction to make, both forms of aggregation are apparently hard to clear by the immune system, as our coincubation experiments with macrophages show (Fig 9, Videos V1-V5). Further studies will be needed to fully understand the implications of media-induced aggregation and antifungal-mediated clustering on clinical treatment. The differences in aggregation types might also explain the varying results linking biofilm formation and aggregation ability in separate studies [17,22,23].

We focused our attention on media-induced aggregation as there is mounting evidence that aggregation of this type is associated with increased biofilm formation and adherence [17]. Given the importance of the cell wall in cell-cell interactions we examined the cell wall ultrastructure comparing *C. auris* cells from different strains grown in RPMI-1640 and SabDex (Fig 5). TEM showed a lack of obvious differences indicating that cell wall mannans are not notably remodelled between aggregated and single cells, supporting the hypothesis of a protein which is not a bulk component of the cell wall mediating media-induced aggregation. We observed moderate, but significant differences in inner cell wall thickness and mannan fibril length between growth condition in some isolates, but these were strain-specific and not associated with a particular clade or the ability to aggregate.

Transcriptomic analysis demonstrated large differences in gene expression between strains of different clades, with 505 DEGs unique to UACa11 (clade I) and 653 DEGs unique to UACa20 (clade III). Of interest to us was that two distinct *ALS* genes were differentially expressed indicating that different ALS-type adhesins might have separate roles [16]. We produced an *als4112* null mutant with some difficulty despite trying multiple techniques, including CRISPR-Cas9-targeted mutation [14]; this might have been aggravated by copy-number variation at the locus [17]. Our *als4112* mutant unfortunately is not a clean deletion, but the region removed does not contain differentially expressed genes identified in our RNA-seq data (Fig S4). Therefore, these genes may not play a role in aggregation and this mutant was thus used for analysis. The results suggest that Als4112 is involved in media-induced aggregation as *als4112* cells formed no aggregates after 24 hours of growth in SabDex. A different mechanism underpins antifungal-induced clustering, as this phenotype was unaffected in *als4112*Δ cells (Figs 2 & 7). This result corroborates the work by Bing *et al.* (2003) where they showed that overexpression of Als4112 induced the aggregation phenotype in non-aggregating strains and that clinical isolates lacking a fully functional Als4112 do not aggregate [17].

The *G. mellonella* infection model has been useful for comparing the pathogenicity of *C. auris* strains [35]. Intriguingly, we show that the choice of growth medium for the inoculum has a significant impact on the outcome (Fig 8A-F). Cells grown in SabDex are more virulent than those grown in RPMI-1640 for all strains tested except one, the non-aggregative clade I strain UACa25. Further testing with a lower starting inoculum of UACa25 (clade I) would rule out a toxic shock phenomenon that could be caused by UACa25 (clade I) having a higher overall virulence compared to other strains (Fig 8A-F). This demonstrates the importance of experimental design and of detailed reporting of experimental conditions to enable a valid comparison across published results. Importantly, when the contribution of aggregation to virulence was directly tested by comparing the wild-type UACa20 isolate to the *als4112*Δ mutant, wild-type infection reduced survival among *G. mellonella* larvae significantly (Fig 8G).

Another difficulty working with aggregation is to correctly determine the number of cells without removing the aggregates from the cell culture before inoculating a host. Therefore, we analysed host-pathogen interaction using macrophages to elucidate how the immune system might deal with *C. auris*. We observed that the uptake of single yeast cells was not impeded in a noticeable way even if aggregates were present (Video V5). Interestingly, macrophages interacted with aggregates but were unable to fully engulf or clear them (Fig 9, Videos V1-V5). We also observed that many macrophages ignored fungal cells which is in line with previous observations [41]. The inability of the immune system to clear aggregates would explain their presence in animal models post infection [18]. We hypothesize that cell aggregates might act as reservoirs in the host during infection, with single cells being cleared by resident macrophages during active infection. However, aggregates might persist for longer times, and cause breakthrough infections when the immune system of the host is weakened or suppressed. Whether this explains the higher virulence of an aggregative wild-type strain compared to a non-aggregative *als4112* mutant requires further experimentation with primary human macrophages and a whole-animal mammalian infection model.

To summarize, we have shown that there are two types of aggregation, canonical strain-inherent aggregation induced by the growth environment, and a different morphology we call “clustering” which is mediated by sub-inhibitory concentrations of echinocandins. We have demonstrated that an ALS-family adhesin, Als4112, is involved in media-induced aggregation, and that aggregates of both types pose a challenge for the immune system and might be a cause of the persistence of *C. auris* systemic infections.

## Materials & Methods

### Yeast strains and culture conditions

All strains used in the study are listed in Table S6. All strains were recovered from storage at −70 °C, grown for 1 day at 37 °C and maintained for up to 1 month on mycological YPD (1% yeast extract (Oxoid Ltd., Basingstoke, UK), 2% mycological peptone (Oxoid Ltd.), 2% D-glucose (Thermo Fisher Scientific, Waltham, MA, USA) and 2% agar (Oxoid Ltd.)) slopes at 4 °C, with fresh cultures spread on mycological YPD plates and incubated at 37°C for 2 days before use. All experiments were performed in Sabouraud dextrose (SabDex) (1% mycological peptone (Oxoid Ltd.), 4% D-glucose (Thermo Fisher Scientific)) or RPMI-1640 (10.4 g/L RPMI-1640 powder without phenol red (Merck KGaA, Darmstadt, Germany), 1.8% D-Glucose (Thermo Fisher Scientific), 0.165 M 3-(N-morpholino)propanesulfonic acid (Melford, Suffolk, UK) adjusted to pH 7 by 1 M NaOH (Sigma-Aldrich, Burlington, MA, USA)). Cultures were grown at 37 °C shaking at 200 rpm. Where plates have been used it is the medium specified with the addition of 2% agar (Becton, Dickinson & Co., Franklin Lakes, NJ, USA).

The antifungal drugs, Caspofungin and Micafungin, were purchased from Merck KGaA and resuspended in DMSO (Merck KGaA). For liquid culture they were added to RPMI- 1640 as prepared above at time of fungal inoculation. For agar plates, the antifungals were added to RPMI-1640 with 2% agar after autoclaving and the media temperature had cooled to 60 °C to avoid degradation of the antifungal activity.

### Determination of Aggregation

Strains were grown for 16 hours at 37 °C shaking at 200 rpm. Cells were harvested by centrifugation (2,500 ×*g*, 5 minutes) after which culture media was removed. Cells where then resuspended in 1× phosphate buffer saline (1× PBS) or ddH2O this was repeated twice to completely remove growth media. Cells finally suspended in 1× PBS or ddH2O were vortexed for 1 minute before microscopic examination to check for presence of aggregates.

### Microscopy

For light microscopy cells were suspended in the indicated solution and 50 µL was added to a glass slide before images were captured on a Zeiss Axioskop 20 microscope with a Axiocam 105 Colour using Zen 2.3 software (version 2.3.69.10000).

For fluorescence microscopy, cells were fixed with 4% methanol-free paraformaldehyde overnight, cells were harvested by centrifugation (2,000 ×*g*, 5 minutes) and resuspended in 1× PBS, this was repeated twice to remove traces of paraformaldehyde.

For cell wall chitin, cells were incubated for 1 hour with 10 µg/mL Calcofluor White M2R (CFW) (Merck KGaA). Cells were harvested by centrifugation (2,000 ×*g*, 5 min) and supernatant removed. Cells were resuspended in 10 µL Vectashield HardSet Antifade Mounting Medium (2B Scientific, Kidlington, UK), which was placed on a microscope glass slide and coverslip placed on top without applying pressure. Images were captured on a Zeiss Imager M2 upright microscope with a Hamamatsu Flash 4 LT camera using Zen 2.3 software (version 2.3.69.1018).

For confocal fluorescent microscopy of chitin and mannan staining, fixed cells were incubated with 1× PBS containing 1% bovine serum albumin (Merck KGaA) for 1 hour on ice. This was removed by centrifugation (2,000 ×*g*, 5 min) and cells were resuspended in 1× PBS containing 10 µg/mL CFW and 25 mg/mL Concanavalin A (Con A) conjugated to Alexa Fluor 488 for 1 hour on ice. Cells were harvested by centrifugation (2,000 ×*g*, 5 min) and resuspended in 1× PBS, this was repeated twice. After the final wash with 1× PBS, cells were spun down again and supernatant was removed, cells were then re-suspended in 10 µL Vectashield HardSet Antifade Mounting Medium and mounted onto microscope slides, a coverslip was applied without pressure. Images were taken as Z- stacks with an image every 0.1 µm with a Ultraview VoX spinning disk confocal microscope using a Hamamatsu C11440-22C camera. Images were processed with Volocity software (Version 6.5.1) selecting images that showed the connection point between cells.

### Galleria mellonella survival assay

Strains were prepared for inoculation by picking a single colony from a SabDex plate and growing the cells for 16 hours in fresh liquid medium (SabDex or RPMI-1640). Medium was removed by centrifugation (2,500 ×*g*, 5 minutes). Cells where then washed twice with ddH2O by centrifugation (2,500 ×*g*, 5 minutes) between each resuspension. Finally, cells were suspended in 1× PBS. Cell suspensions stood for 10 minutes to allow aggregates to sink to the bottom of the suspension and single cells were harvested from the top layer of the solution for counting. Cells were adjusted to give a final inoculum of 50 µL containing 5 × 10^5^ cells in 1× PBS. *G. mellonella* larvae (TruLarv BioSystems Technology, Exeter, UK or UK Waxworms Ltd, Sheffield, UK) were inoculated within two days of receipt. A 50 µL inoculum was given via the last left proleg with a 0.5 ml 29G Micro-Fine U-100 insulin injection unit (BD Medical, Franklin Lakes, NJ, USA).

Larvae were incubated for 6 days at 37 °C. Every 24 hours larvae were examined and deemed dead when they no longer responded to physical stimuli. Each experiment also had a non-injected control group and a control group injected with 1× PBS only. Results were pooled across three independent experiments and statistical analysis was done using a Kaplan-Meier survival plot followed by Log-rank test statistics to determine if differences in survival were significant.

### MIC Testing

Minimum inhibitory concentration (MIC) determination was done using Caspofungin and Micafungin MIC E-test strips (Liofilchem srl, Roseto degli Abruzzi, Italy) following the manufacturer’s guidance with small modifications. Isolates were grown for 24 hours on SabDex agar plates, one colony was picked and dispersed in ddH2O. Onto RPMI-1640 agar plates 2 × 10^6^ cells were spread and allowed to dry for 10 minutes. A MIC test strip was carefully placed onto the agar surface, and the plate was incubated for 24 hours at 37 ^°^C before determining the zone of inhibition and the MIC or MIC90 where appropriate.

### High-pressure freezing sample preparation for transmission electron microscopy (TEM)

Cells were grown for 16 hours in the indicated media. Cells were then harvested by centrifugation (2,500 ×*g*, 5 minutes) and removal of excess cell culture media was performed before fixation and preparation. Fixation and preparation were performed by the Microscopy and Histology Core Facility at the University of Aberdeen following a published protocol [42]. Fixation was done by high-pressure freezing using a Leica Empact 2/RTS high-pressure freezer. Samples were returned to us and we performed imaging using a JEOL 1400 plus transmission electron microscope with an AMT UltraVUE camera. All images were processed, and cell wall thickness was measured with Fiji (ImageJ version 1.53f51) [43].

### Proteinase K assay

Cells were grown for 16 hours in the indicated liquid medium at 37 ^°^C shaking at 200 rpm. Cells were harvested by centrifugation (2,500 ×*g*, 5 minutes), the supernatant was discarded before the pellet was resuspended in 1× PBS, centrifugation and resuspension was repeated twice before aggregates were counted using a haemocytometer. For the assay 1 × 10^6^ cells were added to 50 µL of ddH2O either containing 12.5 µg of proteinase K or not before being incubated for 1 hour at 50 ^°^C, a second ddH2O sample was also prepared and incubated at room temperature for 1 hour to ensure the application of heat did not increase the number of aggregates. After incubation 950 µL 1× PBS was added, and cells were vortexed at high speed for 1 minute. The number of aggregates were counted again with a haemocytometer and the fold-change of aggregates between the unincubated against the incubated samples calculated. Statistical significance was determined by an independent samples t-test.

### Transcriptomics

UACa20 (clade III) and UACa11 (clade I) were grown in the specified media for 16 hours and RNA was extracted for RNA-sequencing (RNA-seq) using the method outlined below, this was repeated three times on independent colonies at different times. After growth for 16 hours cells were harvested by centrifugation (2,500 ×*g*, 5 minutes). Supernatant was discarded and cells resuspended in residual supernatant before immediately proceeding to RNA extraction. RNA was extracted by homogenisation of cells with acid-washed glass beads in 600 µL TRIazole (Invitrogen, Thermo Fisher Scientific, Waltham, MA, USA) per 60 µL culture using a FastPrep-24 5G (MP Biomedicals, Santa Ana, CA, USA) for cell lysis. Glass beads and cell debris were discarded after centrifugation (12,000 ×*g* for 10 minutes at 4 °C) and the supernatant was used for further steps. Addition of 0.2 mL chloroform per 1 mL TRIazole was followed by vigorous shaking for 15 seconds and incubation at room temperature for 2 minutes. Centrifugation (12,000 ×*g* for 15 minutes at 4°C) separated the proteins, DNA, and RNA into 3 phases. RNA in the aqueous layer was removed and subjected to precipitation by addition of 500 µL isopropanol and incubated at room temperature for 10 minutes. Centrifugation (12,000 ×*g* for 10 minutes at 4°C) resulted in a pellet of RNA, which was washed with 600 µL 75% ethanol. Ethanol was removed after centrifugation (6,000 ×*g* for 10 minutes at 4°C) and the RNA pellet was dried for 10 minutes at room temperature. The RNA samples were resuspended in RNase-free water before clean-up treatment with the RNase-Free DNase Set (Qiagen, Hilden, Germany) following the manufacturer’s protocol for off-column clean-up. RNA was then subjected to further clean-up by using the RNeasy Mini Kit (Qiagen) following the manufacturer’s instructions, before final suspension in RNase-free water and storage at −80 °C before sequencing.

Sequencing of mRNA was carried out by the Centre for Genome-Enabled Biology and Medicine (CGEBM) at the University of Aberdeen. Before sequencing External RNA Controls Consortium (ERCC) spike controls were added to samples for assessment of library quality and as estimation of lowest limit of detection [44]. Library preparation was done using Illumina TruSeq Stranded mRNA kit (Illumina Inc., San Diego, CA, USA) and sequenced using the High Output 1×75 kit on the Illumina NextSeq 500 platform producing 75 bp single-end reads. Quality control of the sequencing data was performed using FastQC (version 0.11.8) (https://www.bioinformatics.babraham.ac.uk/projects/fastqc/) with lower-quality reads and adaptor content being removed with TrimGalore! (version 0.6.4) (http://www.bioinformatics.babraham.ac.uk/projects/trim_galore/) with a Phred quality score threshold of 30. ERCC spike controls consist of 92 synthesized transcripts that range both in length and concentration. Two mixes (Mix 1 and Mix 2) are provided with differing transcript concentration ratios of 4:1, 1:1, 1:1.5, and 2:1 to allow for assessment of detection of differential expression. One of these two mixes was randomly added to each of the samples. ERCC reads were removed by aligning all reads to the ERCC reference genome using HISAT2 (version 2.1.0) [45] with unmapped *C. auris* reads being kept for alignment. Illumina-sequencing produced between ∼25 million and ∼30 million reads per sample (Table S7).

The *C. auris* B11221 reference genome was downloaded from NCBI (https://www.ncbi.nlm.nih.gov/genome/38761?genome_assembly_id=678645) [46] along with the equivalent annotation file and prepared for use with the alignment software HISAT2 (version 2.1.0) [45]. Reads were aligned to this prepared reference using HISAT2 (version 2.1.0) [45] with the parameter for stranded library preparation used. SAMtools (version 1.9) [47] was used to process the alignments and reads were counted at gene locations using featureCounts (part of the sub-read version 1.6.2 package) [48] utilizing the parameter to split multi-mapped reads as a fraction across all genes that they align to, and the parameter for stranded analysis. 83.66% to 88.75% of reads aligned to coding regions and after quality control and removal of low-count genes a total of 5,431 genes remained for analysis.

edgeR (version 3.30.3) [49] was used to detect which genes had a significant differential change in expression. All genes that had a CPM (count per million) value of more than one in three or more samples were kept for analysis, and all other genes were removed as low-count genes. Differential expression analysis was performed via pairwise comparisons between each sample group. To obtain the B8441 gene names, the gene DNA sequence for the B11221 strain was obtained and compared to the B8441 gene sequences using BlastN [50].

### *als4112* mutant generation

All genomic DNA was prepared as described previously [12]. All PCRs were done using VeriFi Hot Start Mix (PCR Biosystems Ltd., London, UK), with the exception of transformant screening which was done with PCRBIO Taq Mix Red (PCR Biosystems Ltd.). The deletion mutant was generated by a previously described method of homology-directed repair and lithium-acetate transformation [51] using a nourseothricin marker. Briefly, the *CaNAT1* resistance marker was amplified from plasmid pALo218 using primers oUA315 and oUA316 (Table S8) [12]. The *ALS4112* gene sequence and flanking sequences 2 kb upstream and downstream were obtained from NCBI (Accession XP_028889036.1) to design primers, 1 kb upstream and 1 kb downstream of *ALS4112* was amplified with primers oUA989 and oUA990, and oUA991 and oUA992 respectively, from UACa20 genomic DNA (Table S8). The deletion cassette was assembled using fusion PCR and primers oUA993 and oUA994 (Table S8) to amplify the final deletion cassette which was confirmed by gel electrophoresis. UACa20 (clade III) was grown overnight at 30 °C with shaking before being diluted 1:100 in fresh bacteriological YPD (1% yeast extract (Oxoid Ltd.), 2% bacteriological peptone (Oxoid Ltd.), 2% D-glucose (Thermo Fisher Scientific) and 2% agar (Oxoid Ltd.)) and grown for a further 4 hours. Cells were then harvested by centrifugation for 2 minutes at 1,000 ×*g* and resuspended in ddH2O. This step was repeated twice before cells were suspended in 0.1 M lithium acetate and were centrifuged again for 1 minute at 1,000 ×*g* before being suspended in 0.1 M lithium acetate. To this 5 µg of the deletion cassette in 50 µL of 50% PEG-3350, 1 M lithium acetate and 10 mg/mL denatured herring sperm DNA (Thermo Fisher Scientific) was added and incubated overnight shaking at 30 °C. Heat shock was applied in a heat block the next day at 44 °C for 15 minutes, and cells were resuspended in bacteriological YPD and incubated shaking at 30 °C for 3 hours to recover. After recovery cells were plated on bacteriological YPD plates containing 200 µg/mL nourseothricin (clonNAT; Werner BioAgents GmbH, Jena, Germany) and incubated at 30 °C until colonies appeared (usually 2 days). Transformants were screened for *als4112* deletion with primers oUA987 and oUA988 (Table S8), and UACa20 genomic DNA was used as a positive control.

### Inverse PCR

Genomic DNA from the *als4112* null mutant was prepared as described previously [12]. Inverse PCR design is outlined in Fig S4. For the restriction digest 1 µg of genomic DNA from UACa20 and *als4112*Δ was incubated for 1 hour at 37 °C with the restriction endonuclease BamHI-HF (New England Biolabs (NEB), Ipswich, MA, USA) following the manufacturer’s instructions. The enzyme was removed with the Monarch PCR & DNA Cleanup Kit (NEB) following the manufacturer’s instructions. For the ligation reaction 8 µL of the digest was treated with T4 Ligase (NEB) for 16 hours at 16 °C, before stopping the reaction at 65 °C for 10 minutes. PCR was done using VeriFi Hot Start Mix (PCR Biosystems Ltd.) to amplify DNA for sequencing with primers oUA995 and oUA996 (Table S8) for upstream region boundary of the deletion cassette and primers oUA997 and oUA998 (Table S8) were used for the downstream boundary of the deletion cassette, amplification of the *als4112*Δ DNA but not UACa20 DNA was confirmed by gel electrophoresis. DNA was sent for Sanger sequencing (Eurofins Scientific, Luxembourg, Luxembourg) with primers oUA999 for upstream and oUA1000 for downstream sequencing (Table S8), and results were compared using BLAST against the *C. auris* reference genome (NCBI Cand_auris_B11221_V1) to determine boundaries of the deletion cassette.

### THP-1 Macrophage Differentiation

The human monocyte cell line THP-1 (MerckKGaA) were maintained at 2 × 10^5^ cells/mL in RPMI-1640 containing 10% fetal bovine serum (FBS) (Thermo Fisher Scientific), 200 U/mL penicillin/streptomycin (Thermo Fisher Scientific), and 2 mM GlutaMAX (Thermo Fisher Scientific) at 37 °C in 5% CO2. Macrophages were differentiated with 200 nM phorbol 12-myristate 13-acetate (PMA) (Merck KGaA) following a previously described method with slight modifications [52]. Briefly 1 × 10^5^ cells/ml were incubated in RPMI- 1640 with 200 nM PMA, 10% FBS, 200 U/mL penicillin/streptomycin, and 2 mM GlutaMAX at 37 °C in 5% CO2 for 3 days. PMA-containing media and non-adherent cells were removed by washing twice with 1× PBS. THP-1 macrophages were then allowed to rest for 3 days in RPMI-1640 with 10% FBS, 200 U/mL penicillin/streptomycin, and 2 mM GlutaMAX at 37 °C in 5% CO2. Media was removed and THP-1 macrophages were lifted by scraping. 2 × 10^5^ THP-1 macrophages were added to each well of an 18-well slide (ibidi, Gräfelfing, Germany) in RPMI-1640 supplemented with 10% FBS, 200 U/mL penicillin/streptomycin, and 2 mM GlutaMAX at 37 °C in 5% CO2 and allowed to rest for a further 24 hours before being used for experimentation.

### Live-Cell Imaging

Fungal cells were grown for 24 hours in desired media and cells were harvested by centrifugation 2,000 ×*g* for 5 minutes and resuspended in ddH2O, this was repeated twice before cells were counted with a haemocytometer. Medium was removed from THP-1 macrophages and replaced with CO2-independent medium (Fisher Scientific) containing fungal cells at MOI 1:3 and 0.02 µg/µL propidium iodide (Fisher Scientific). Images were taken every 3 min for 2 hours on an Ultraview VoX spinning disk confocal microscope using Volocity software (Version 6.5.1).

### Statistics

All statistics were performed in SPSS Statistics for Windows (version 28.0) (IBM Corp., Armonk, NY, USA) except for log-rank tests of *G. mellonella* survival data which were performed in R (version Rx64, 4.3.0) (http://www.r-project.org/).

## Acknowledgments

We are grateful to Rhys Farrer (MRC CMM, University of Exeter) for guidance on bioinformatic analysis, to Louise Walker (University of Aberdeen) for advice on macrophage work, and to Neil A. R. Gow (MRC CMM, University of Exeter) and Dhara Malavia (MRC CMM, University of Exeter) for discussing their unpublished results. We thank Gillian Milne (Microscopy & Histology Facility, University of Aberdeen) for technical assistance with electron microscopy. *Candida auris* strains were kindly provided by Anuradha Chowdhary (Vallabhbhai Patel Chest Institute, University of Delhi, Delhi, India), Public Health England (PHE) (Bristol, UK), and the Mycotic Diseases Branch of the Centers for Disease Control and Prevention (CDC) (Atlanta, GA, USA).

## Author Summary

*Candida auris* is a single-celled fungus, a yeast, that can cause severe infections in hospital patients. This fungus is difficult to treat because it is resistant to many antifungal drugs. Therefore, to understand the processes that enhance the virulence of this yeast with a view to developing new treatments. Previous studies have found that *C. auris* can form aggregates, or clumps of cells, which may play a role in how the fungus infects people. In this study, we identified two different types of aggregation in *C. auris*, one triggered by antifungal treatment, and another controlled by growth conditions. This discovery allowed us to study aggregate formation in the same genetic background. In doing so, we found that a certain protein, an ALS-family adhesin, is involved in the aggregation process. Surprisingly, we also discovered that aggregates may promote infection by making it harder for the immune system to clear the yeast. This new understanding could help researchers develop better ways to fight *C. auris* infections.

