## Supplementary Figures & Tables for "*Candida auris* undergoes adhesin-dependent and -independent cellular aggregation"

**Table S1.** $MIC_{90}$ of Caspofungin (CSP) and Micafungin (MFG), $MIC_{90}$ as identified by E-test strips

| Strain | CSP (mg/L) $MIC_{90}$ | MFG (mg/L) $MIC_{90}$ |
|---|---|---|
| UACa11 | 12 | 0.094* |
| UACa25 | 1 | 0.094* |
| UACa10 | 0.175 | 0.094 |
| UACa20 | 0.38 | 0.064 |

*Actual MIC with no growth above indicated concentration.

**Table S2.** Genes upregulated in both *C. auris* UACa20 and UACa11 during growth in SabDex but with a greater than log2FC difference between strains

| Gene | UACa20 log2FC | UACa11 log2FC | Potential homolog in *C. albicans* |
|---|---|---|---|
| CJI97_004514 | 9.49 | 7.42 | *THI13* |
| CJI97_003172 | 7.55 | 9.66 | *THI4* |
| CJI97_002670 | 4.24 | 2.07 | C3_00990C_A |
| CJI97_000357 | 4.19 | 1.14 | - |
| CJI97_000368 | 3.64 | 1.61 | C3_03440C_A |
| CJI97_001224 | 2.11 | 4.55 | *RBT7* |
| CJI97_001401 | 1.95 | 4.08 | *THI6* |
| CJI97_004176 | 1.66 | 3.85 | *MDR1* |

**Table S3.** DEGs expressed in opposite direction between strains UACa20 and UACa11

| Gene | UACa20 log2FC | UACa11 log2FC | Potential homolog in *C. albicans* |
|---|---|---|---|
| CJI97_001729 | 3.75 | -1.45 | CR_01630C_A |
| CJI97_004216 | 1.67 | -1.01 | *HBR1* |
| CJI97_000418 | 1.53 | -1.23 | *CFL4* |
| CJI97_004611 | 1.53 | -2.70 | *CDA2* |
| CJI97_001311 | 1.52 | -2.84 | C6_02660C_A |
| CJI97_000125 | 1.36 | -1.10 | *TSR1* |
| CJI97_005371 | 1.28 | -2.36 | *SAP5* |
| CJI97_004564 | 1.21 | -1.90 | *SAP8* |
| CJI97_004013 | -1.21 | 1.36 | *PLB4.5* |
| CJI97_001240 | -1.22 | 1.23 | C4_07040W_A |
| CJI97_004556 | -1.78 | 3.55 | *PRD1* |
| CJI97_005598 | -2.42 | 1.37 | C2_06350C_A |
| CJI97_004563 | -2.66 | 1.67 | *DUR3* |

**Table S4.** Top 10 upregulated DEGs during growth in SabDex unique to *C. auris* strain UACa11

| Gene | log2FC | Potential homolog in *C. albicans* |
|---|---|---|
| CJI97_002126 | 3.94 | *ALS4* |
| CJI97_004172 | 3.71 | *RBR3* |
| CJI97_002948 | 3.27 | C1_10310W_A |
| CJI97_004914 | 2.90 | *OPT3* |
| CJI97_002178 | 2.88 | *CYB2* |
| CJI97_004170 | 2.87 | - |
| CJI97_004560 | 2.75 | *HYR3* |
| CJI97_004890 | 2.71 | *OPT1* |
| CJI97_004892 | 2.59 | *OPT3* |
| CJI97_005489 | 2.55 | *TRA1* |

**Table S5.** Top 10 upregulated DEGs during growth in SabDex unique to *C. auris* strain UACa20

| Gene | log2FC | Potential homolog in *C. albicans* |
|---|---|---|
| CJI97_000210 | 3.70 | C3_02040C_A |
| CJI97_004175 | 3.67 | *ALS4* |
| CJI97_003074 | 3.50 | - |
| CJI97_003045 | 3.29 | *VHT1* |
| CJI97_004212 | 3.04 | *FAD3* |
| CJI97_000710 | 2.90 | C3_04450C_A |
| CJI97_002719 | 2.71 | *AGP2* |
| CJI97_003695 | 2.70 | C2_04080W_A |
| CJI97_003085 | 2.62 | *SRP40* |
| CJI97_000246 | 2.57 | C1_10970W_A |

**Table S6.** List of *Candida auris* strains

| Strain # | Isolate Name | Clade | Source/Reference | Aggregative ability |
|---|---|---|---|---|
| **UACa6** | NCPF8980#9 | III | PHE, E. Johnson | Yes |
| **UACa10** | NCPF13005#95 | III | PHE, E. Johnson | Yes |
| **UACa11** | VPCI479/P/13 | I | [1] | No |
| **UACa25** | B11098 | I | CDC [3] | No |
| **UACa20** | B11221 | III | CDC [3] | Yes |
| **UACa22** | B11244 | IV | CDC [3] | Yes |
| **UACa83** | CBS10913T | II | [2] | No |
| **UACa177** | *als4112Δ* | III | Derivative of UACa20 | No |

**Table S7.** Sequencing quality report for transcriptomic data

| **Strain** | UACa20 | UACa20 | UACa11 | UACa11 |
|---|---|---|---|---|
| **Condition** | SabDex (Agg) | RPMI-1640 (Non-Agg) | SabDex (Agg) | RPMI-1640 (Non-Agg) |
| **Raw Reads** | **29,430,966** | **27,484,339** | **27,470217** | **26,102,812** |
| **Filtered Reads** | **29,013,091** | **26,502,460** | **27,195,036** | **25,870,831** |
| **Percentage (%) of filtered reads** | | | | |
| aligned to ERCC | 0.54 (0.28) | 0.51 (0.21) | 0.45 (0.17) | 0.47 (0.18) |
| aligned to B11221 genome | 98.76 (0.30) | 98.59 (0.33) | 97.79 (0.22) | 97.25 (0.26) |
| aligned as multimappers | 1.29 (0.04) | 2.35 (0.65) | 1.75 (0.18) | 2.61 (0.66) |
| counted as genes | 88.49 (0.23) | 85.24 (0.89) | 86.48 (0.27) | 84.55 (0.83) |

**Reads are the average of the three biological repeats rounded to the nearest whole number. Percentages of filtered reads aligned are given as means with standard deviation in parentheses.**

**Table S8.** Oligonucleotide primer list

| Primer | Sequence (5’-3’) |
|---|---|
| oUA315 | TCGTACGCTGCAGGTCGACGGTCGACACTGGATGGCGG |
| oUA316 | CGCGCCTTAATTAACCCGGGATCAAGCTTGCCTCGTCC |
| oUA989 | CCTTATTGCACCCGTGG |
| oUA990 | CGATACTAACGCCGCCATCCAGTGTCGACAGGAAAGTGGAAATTGTGGG |
| oUA991 | GGCGGGGACGAGGCAAGCTTGATCTATGTCTCACACCAAGG |
| oUA992 | GCATTCAAGATGAGATTATTG |
| oUA993 | GCACACTCGACAAGTCTTAGGC |
| oUA994 | CAACAATTTAATATCATAGGAGCTCACG |
| oUA987 | GGCGAAACTGTCACTGTTGT |
| oUA988 | GGTGGCTCAGTGAAGATCCT |
| oUA995 | CAATAGCTTCAGCATCACCTGG |
| oUA996 | CTTTGATTATCAAAGGTTCTCGCTTAGCC |
| oUA997 | GTCTCGTTGCTTAACTGCTGC |
| oUA998 | GGATACAGTTCTCACATCACATCC |
| oUA999 | CCACCTCCCTCGCCAGAC |
| oUA1000 | GCCAGTCGTGATGTGTGC |

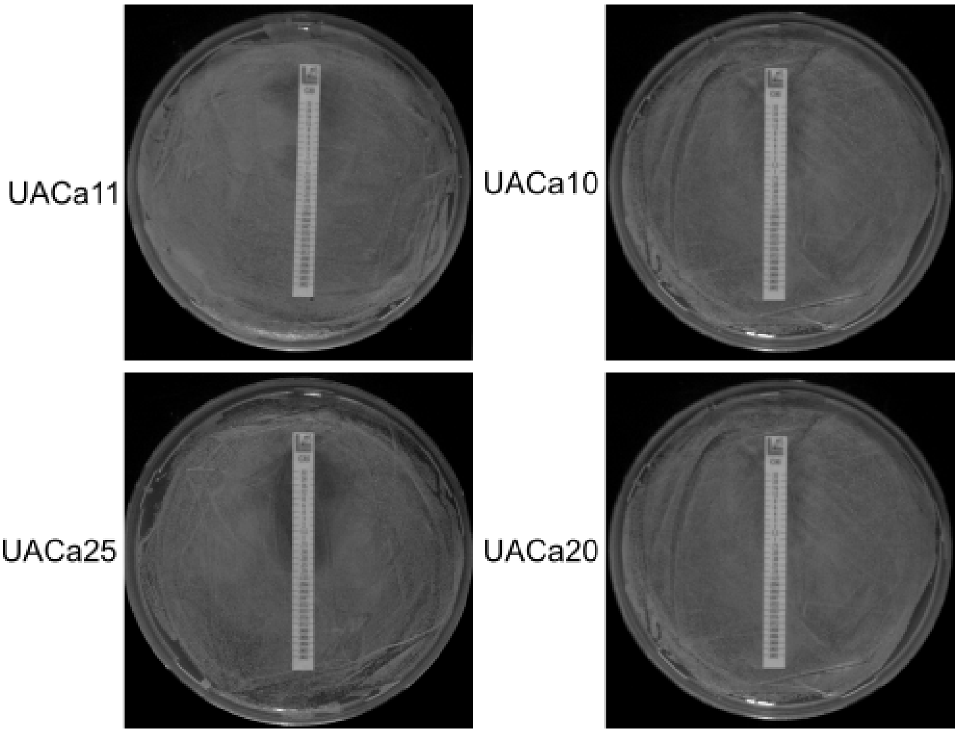


**Fig S1. Caspofungin E-test was used to determine the MIC of indicated strains.** All strains had growth up to the 32 mg/L mark along the E-test strip. The $MIC_{90}$ was determined from where growth was decreased, for UACa11 this was 12 mg/L, for UACa25 1 mg/L, for UACa10 0.175 mg/L, and for UACa20 0.38 mg/L.

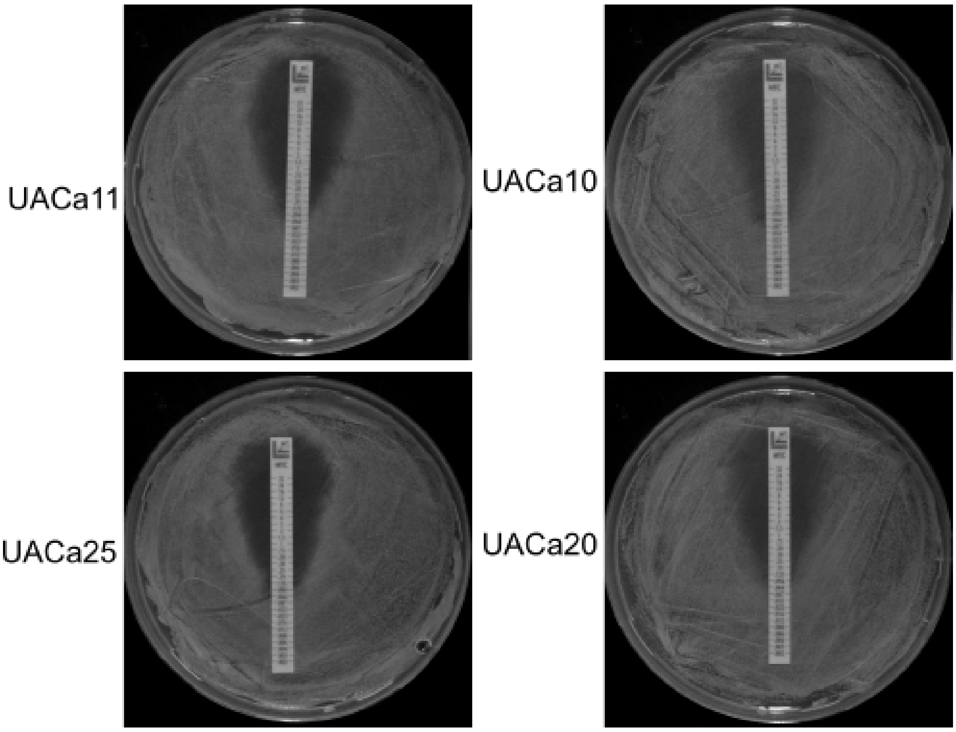


**Fig S2. Micafungin MIC E-test was used to determine the MIC of indicated strains.** UACa11 and UACa25 had clear zones of inhibition and the MIC was 0.094 mg/L for both. UACa10 and UACa20 did not show a clear zone of inhibition, but a zone with reduced growth, the $MIC_{90}$ were estimated to be 0.094 mg/L and 0.064 mg/L, respectively.

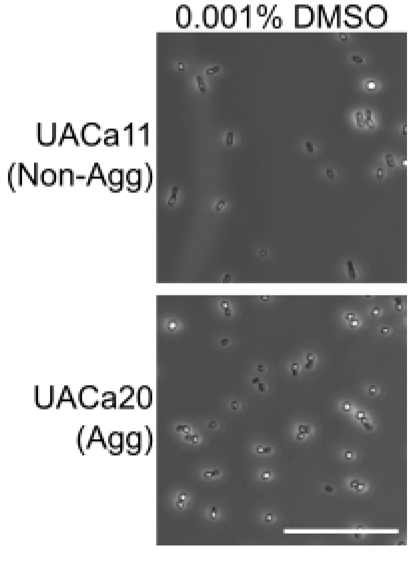


**Fig S3. Growth in RPMI-1640 with 0.001% DMSO does not change the phenotype of cells.** UACa11 and UACa20 grown in RPMI-1640 with 0.001% DMSO as a control to ensure that the antifungal carrier does not cause growth defects. Scale bar represent 50 µm.

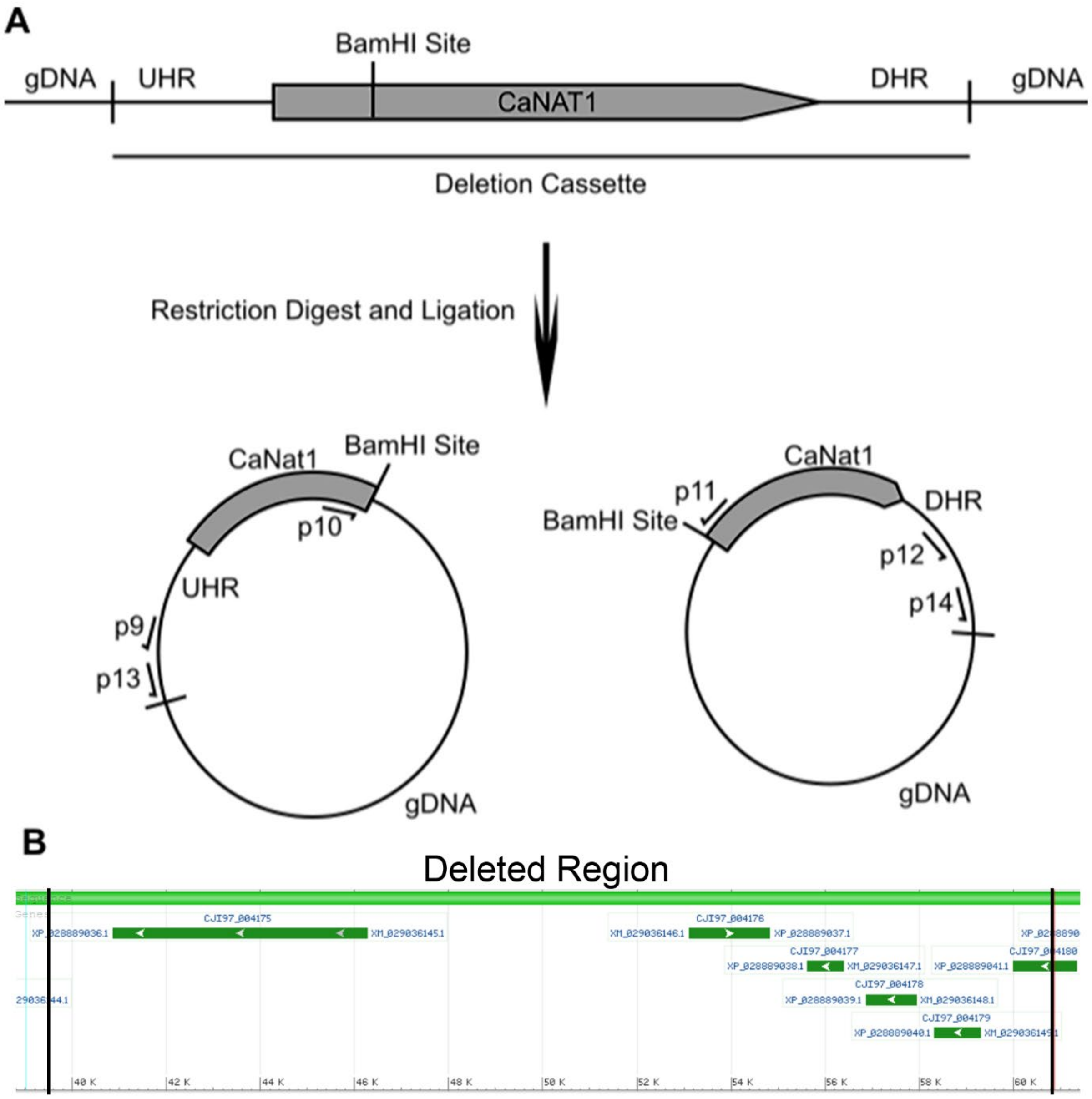


**Fig S4. Inverse PCR to identify *CaNAT1* deletion cassette integration location.** (**A**) Genomic DNA from the *als4112*Δ mutant was digested with BamHI to split the *CaNAT1* marker locus and fragment the genomic DNA, ligation followed to form circular DNA that could be amplified with PCR. Primers designed to amplify parts of *CaNAT1* and the upstream homology region (UHR) or parts of *CaNAT1* and the downstream homology region (DHR) of the deletion cassette were used to generate PCR products which were subjected to Sanger sequencing. (**B**) Black lines indicate the boundaries of the *als4112* deletion as determined by BLAST searches of the sequences from the inverse PCR against the B11221 genome assembly on NCBI.

### Video Legends

**Video V1. UACa20 cells grown in RPMI-1640 interacting with THP-1 derived macrophages.** THP-1 derived macrophages were co-incubated with fungal cells grown in RPMI-1640 at a MOI 1:3 (Differential Inference Contrast overlaid with red fluorescence channel). Fungal cells are engulfed without complications. Interactions were recorded over 2 hours with 1 image taken every 3 minutes. The red fluorescence channel detects propidium iodide staining which was included to highlight dead macrophages and yeast cells. Scale bar represents 15 µm.

**Video V2. UACa20 cells grown in RPMI-1640 with 0.0001% DMSO interacting with THP-1 derived macrophages.** THP-1 derived macrophages were co-incubated with fungal cells grown in RPMI-1640 with 0.0001% DMSO at a MOI 1:3 (Differential Inference Contrast overlaid with red fluorescence channel). Fungal cells are engulfed without complications. Interactions were recorded over 2 hours with 1 image taken every 3 minutes. The red fluorescence channel detects propidium iodide staining which was included to highlight dead macrophages and yeast cells. Scale bar represents 15 µm.

**Video V3. UACa20 aggregates grown in SabDex interacting with THP-1 derived macrophages.** THP-1 derived macrophages were co-incubated with fungal cells grown in SabDex at a MOI 1:3 (Differential Inference Contrast overlaid with red fluorescence channel). Individual macrophages fail to engulf aggregates. Interactions were recorded over 2 hours with 1 image taken every 3 minutes. The red fluorescence channel detects propidium iodide staining which was included to highlight dead macrophages and yeast cells. Scale bar represents 15 µm.

**Video V4. UACa20 clusters grown in RPMI-1640 with 0.075 mg/L MFG interacting with THP-1 derived macrophages.** THP-1 derived macrophages were co-incubated with fungal cells grown in RPMI-1640 containing 0.075 mg/L MFG at a MOI 1:3 (Differential Inference Contrast overlaid with red fluorescence channel). Here, macrophages fail to engulf clusters of cells. Interactions were recorded over 2 hours with 1 image taken every 3 minutes. The red fluorescence channel detects propidium iodide staining which was included to highlight dead macrophages and yeast cells. Scale bar represents 15 µm.

**Video V5. Single cells of UACa20 grown in SabDex interacting with THP-1 derived macrophages.** THP-1 derived macrophages were co-incubated with fungal cells grown in SabDex at a MOI 1:3 (Differential Inference Contrast overlaid with red fluorescence channel). Interactions were recorded over 2 hours with 1 image taken every 3 minutes. Interactions with single cells grown in SabDex are the same as the cells grown in RPMI-1640 (see Videos V1 and 2 for comparison). The red fluorescence channel detects propidium iodide staining which was included to highlight dead macrophages and yeast cells. Scale bar represents 15 µm.
